# Investigating resting brain perfusion abnormalities and target-engagement by intranasal oxytocin in women with bulimia nervosa and binge-eating disorder and healthy controls

**DOI:** 10.1101/2020.01.26.920090

**Authors:** Daniel Martins, Monica Leslie, Sarah Rodan, Fernando Zelaya, Janet Treasure, Yannis Paloyelis

## Abstract

Advances in the treatment of bulimia nervosa and binge eating disorder (BN/BED) have been marred by our limited understanding of the underpinning neurobiology. Here we measured regional cerebral blood flow (rCBF) to map resting perfusion abnormalities in women with BN/BED compared to healthy controls and investigate if intranasal oxytocin (OT), proposed as a potential treatment, can restore perfusion in disorder-related brain circuits. Twenty-four women with BN/BED and 23 healthy women participated in a randomised, double-blind, crossover, placebo-controlled study. We used arterial spin labelling MRI to measure rCBF and the effects of an acute dose of intranasal OT (40IU) or placebo over 18-26 minutes post-dosing, as we have previously shown robust OT-induced changes in resting rCBF in men 15-36min post-dosing. We tested for effects of treatment, diagnosis and their interaction on extracted rCBF values in anatomical regions-of-interest previously implicated in BN/BED by other neuroimaging modalities, and conducted exploratory whole-brain analyses to investigate previously unidentified brain regions. We demonstrated that women with BN/BED presented increased resting rCBF in key neural circuits previously implicated in BN/BED pathophysiology, namely the medial prefrontal and orbitofrontal cortices, anterior cingulate gyrus, posterior insula and middle/inferior temporal gyri bilaterally. Hyperperfusion in these areas specifically correlated with eating symptoms severity in patients. Our data did not support a normalizing effect of 40IU intranasal OT on perfusion abnormalities in these patients 18-26 minutes post-dosing. Our findings enhance our understanding of resting brain abnormalities in BN/BED and identify resting rCBF as a non-invasive potential biomarker for disease-related changes and treatment monitoring.

## INTRODUCTION

Bulimia nervosa (BN) and binge eating disorder (BED) are psychiatric disorders characterized by recurrent binge eating^1^. In BN, binge eating is accompanied by excessive compensatory behaviours, which are absent in BED^2^. Since recovery rates after standard treatment remain poor in both disorders^3^, developing new effective treatments that address binge eating are needed; however, progress is impeded by our limited understanding of the neurobiology of BN/BED.

BN/BED have been conceptualized as impulsive/compulsive eating disorders^4^ with altered reward sensitivity^5^ and food-related attentional biases^6^. Consistent with this model of BN/BED, neuroimaging studies in these patients have highlighted changes in the function, structure and neurochemistry in brain areas involved in incentive processing (such as the orbitofrontal cortex, ventral tegmental area(VTA), substantia nigra(SN), nucleus accumbens and the amygdala), inhibitory control (such as the medial prefrontal cortex (PFC) and the anterior cingulate (ACG)) and habitual behaviour (such as the dorsal striatum) (see^7, 8^ for detailed reviews). Most of these studies focused on morphological differences between BN/BED patients and healthy controls. Some studies investigated functional abnormalities using task-based BOLD functional magnetic resonance imaging (fMRI), allowing for inferences that are restricted to the neural circuits employed by the specific tasks. Other studies investigated alterations in spontaneous fluctuations of the BOLD signal at rest, suggesting aberrant functional connectivity in brain networks involved in salience attribution, self-referential processing and cognitive control in people with BN/BED^9–13^). However, BOLD fMRI (whether focusing on fluctuations at rest or task-evoked changes) typically yields only relative metrics and is prone to several artefacts, including motion, low-frequency physiological noise and baseline drift^14^. Three other studies, all with small sample sizes, used Single Photon Emission Computerized Tomography (SPECT) to measure regional cerebral blood flow (rCBF), which provides a proxy of brain metabolism and neural activity^15^. These studies reported rCBF increases in the temporal lobes and/or the frontal cortex of patients with BN and BED in response to anticipation or exposure to food/body shape-related stimuli^16–18^. However, SPECT is a high cost technique that requires the injection of a radionuclide tracer, making it suboptimal as a routine screening tool in non-specialized centres, or for evaluating short-term responses^19^.

Recently, arterial spin labelling (ASL) MRI has allowed the non-invasive quantification of rCBF with high reproducibility and spatial resolution^20^. ASL is a quantitative technique and has been used widely to quantify changes in brain physiology associated with various neuropsychiatric disorders^21^ (*e.g.,* Autism Spectrum Disorder^22^, Schizophrenia^23^, Depression^24^) and in response to pharmacological treatments^25, 26^. Hence ASL provides a non-invasive and widely available tool to investigate abnormalities in resting brain physiology of BN/BED and the effects of potential treatments.

One such potential treatment is intranasal oxytocin (OT)^27–29^. Support for the implication of OT in binge eating comes from animal^30^ and human studies^31, 32^ demonstrating that OT suppresses eating, including hedonic eating in men^33^. Some small clinical studies have also shown that intranasal OT (40IU) reduces caloric intake^34^ and decreases vigilance towards angry faces^35^ in women with BN. Additional evidence for the involvement of the OT system in dysregulated eating comes from studies associating polymorphisms in the OT receptor gene with binge-purge tendencies in women (rs53576/rs2254298)^36^, bulimia (rs53576)^37^ and overeating (rs2268493/rs2268494)^38^. However, it is yet not clear which brain circuits intranasal OT targets in BN/BED patients.

In this study, we investigate abnormalities in resting brain physiology in women with BN/BED using ASL. Studying the brain at rest allows us to investigate baseline alterations in brain physiology that are not restrained to the specific neural circuits engaged by tasks. We further investigate whether an acute dose of 40IU of intranasal OT can restore alterations in resting brain physiology in BN/BED 18-26 minutes post-doing. We have previously shown that ASL captures OT-induced changes as early as 15-36 mins post-dosing in resting rCBF after a single acute intranasal administration (40 IU) in both healthy men^39, 40^ and men at clinical high-risk for psychosis (CHR-P)^41^ (earlier time intervals have simply never been studied). We specifically address two questions: (1) Do women with BN/BED, compared to healthy women, present alterations in resting brain perfusion, as measured using ASL MRI? (2) Can 40IU intranasal OT restore or attenuate these resting rCBF alterations in women with BN/BED 18-26 minutes post-doing?

## METHODS

### Participants

We recruited 25 women meeting the DSM-5 criteria for either BN (N=20) or BED (N=5) and 27 women with no current or prior eating disorder. We excluded three participants (2 healthy controls and 1 BN patient) due to a large discrepancy in the time post-dosing that we sampled rCBF in the OT and placebo visits (it exceeded the duration of one rCBF scan, i.e. >8 mins). We further excluded two healthy women due to corruption of the data files. Our final sample included 23 healthy and 24 women with BN/BED. Ethical approval for the study was granted by the London – Camberwell St Giles Research Ethics Committee (Reference: 14/LO/2115). See supplementary material for further details regarding inclusion/exclusion criteria. Our previous work has demonstrated that N=16 per group is sufficient to quantify OT-induced rCBF changes in men using a between-subjects^42^ or within-subjects design^40^.

### Study Design and Procedure

We employed a double-blind, placebo-controlled cross-over design. Participants visited our centre on three occasions: one screening and two experimental visits. During screening, we measured height, weight, and collected basic demographics. Participants then completed the Eating Disorder Examination – Questionnaire Version^43^ and Depression, Anxiety, and Stress Scales^44^ online in their own time between the screening and the first experiment visits. Both questionnaires have been validated for online application^45, 46^. The experimental visits were conducted two days apart, ensuring participants were tested in the same phase of the oestrous cycle for both treatment conditions (OT, placebo). Each participant was asked to report the first day of their last menstrual period and any hormonal contraception they were currently taking. Participants were asked to eat 2.5 hours prior to each experimental visit to control for baseline hunger. All participants were tested at approximately the same time in the early evening (5-7 pm) for both the OT and placebo treatments, to minimise potential circadian variability in resting brain activity ^47^ or OT levels^48^. Fifty minutes after arrival, participants self-administered 40IU intranasal OT (Syntocinon, 40IU/ml; Novartis, Basel, Switzerland) in 10 puffs, one puff every 30s, each puff containing 0.1ml Syntocinon (4IU) or placebo (same excipients as Syntocinon except for OT) and alternating between nostrils, over a period of 5 minutes. Participants were randomly allocated to a treatment order (OT/placebo or placebo/OT). After drug administration, participants were guided to the MRI scanner, where a single pulsed continuous ASL scan (8:20 minutes) was acquired 18-26 minutes (± 4 mins) post-dosing. Our choice of OT dose and post-dosing interval was driven by our previous work, whereby we have shown that 40IU intranasal OT induce robust changes in rCBF in both healthy men (as early as 15-32 mins post-dosing)^40, 42^ and men at CHR-P (at 22-36 mins)^41^. Furthermore, previous clinical studies have shown beneficial effects of a dose of 40IU intranasal OT in women with BN^34, 35^.

### MRI data acquisition

We used a 3D pseudo-continuous ASL (3D-pCASL) sequence to measure changes in regional Cerebral Blood Flow (rCBF) over 18-26 min post-dosing. Participants were instructed to lie still and maintain their gaze on a centrally placed fixation cross during scanning. Labelling of arterial blood was achieved with a 1525 ms train of Hanning shaped RF pulses in the presence of a net magnetic field gradient along the flow direction (the z-axis of the magnet). After a post-labelling delay of 2025ms, a whole brain volume was read using a 3D inter-leaved “stack-of-spirals” Fast Spin Echo readout^49^, consisting of 8 interleaved spiral arms in the in-plane direction, with 512 points per spiral interleave. TE was 11.088 ms and TR was 5135 ms. 56 slice-partitions of 3mm thickness were defined in the 3D readout. The in-plane FOV was 240×240 mm. The spiral sampling of k-space was re-gridded to a rectangular matrix with an approximate in-plane resolution of 3.6 mm. Each sequence used 5 control-label pairs. Individual CBF maps were computed for each of the perfusion weighted difference images derived from every control-label (C-L) pair, by scaling the difference images against a proton density image acquired at the end of the sequence, using identical readout parameters. This computation was done according to the formula suggested in the recent ASL consensus article^50^. The sequence uses four background suppression pulses to minimise static tissue signal at the time of image acquisition. We acquired eight 3D-pCASL sequences, with the duration of the entire acquisition time of each sequence being 8:20 min.

A 3D high-spatial-resolution, Magnetisation Prepared Rapid Acquisition (3D MPRAGE) T1-weighted scan was also acquired (field of view of 270mm, TR/TE/TI = 7.328/3.024/400ms). The final resolution of the T1-weighted image was 1.1 x 1.1 x 1.2 mm.

### MRI data pre-processing

A multi-step approach was performed for the spatial normalization of the CBF maps to Montreal Neurological Institute (MNI) space: (1) co-registration of the proton density image from each sequence to the participant’s T1-image after resetting the origin of both images to the anterior commissure. The transformation matrix of this co-registration step was then applied to the CBF map, to transform the CBF map to the space of the T1-image; (2) unified segmentation of the T1 image; (3) elimination of extra-cerebral signal from the CBF map, by multiplication of the “brain only” binary mask obtained in step[2], with each co-registered CBF map; (4) normalization of the subject’s T1 image and the skull-stripped CBF maps to the MNI152 space using the normalisation parameters obtained in step[2]. Finally, we spatially smoothed each normalized CBF map using an 8-mm Gaussian smoothing kernel. All of these steps were implemented using the ASAP (Automatic Software for ASL processing) toolbox (version 2.0)^51^. The resulting smoothed CBF maps were then entered into Statistical Parametric Mapping (SPM) 12 (http://www.fil.ion.ucl.ac.uk/spm/software/spm12/) for group-level statistical analysis, as described below.

## Statistical analyses

### Diagnosis, treatment and diagnosis x treatment effects on resting rCBF

To investigate the effects of diagnosis and treatment on global CBF, we first extracted mean global CBF values with an explicit binary mask for grey-matter using the fslmeants command (FSL suite, http://www.fmrib.ox.a.c.uk/fsl). The binary mask was derived from a standard T1-based probabilistic map of grey-matter distribution by thresholding all voxels with a probability >.20. We tested for main effects of treatment or diagnosis, and for the treatment x diagnosis interaction on global grey-matter CBF signal using mixed analysis of variance and the Greenhouse-Geisser correction against violations of sphericity.

We then tested the effects of diagnosis, treatment and diagnosis x treatment on mean rCBF values extracted using *fslmeants* from 14 regions-of-interest (ROIs) corresponding to anatomical areas shown to be affected in BN/BED in previous structural, functional or neurochemical studies^7, 8^. A detailed description of these ROIs can be found in Table 2 and Fig. S1. The VTA, SN and hypothalamus masks were retrieved from a recently published high-resolution probabilistic atlas of human subcortical brain nuclei^52^. The orbitofrontal cortex (OFC) mask included the areas 14 and 11m, and the medial PFC included area FPm from the connectivity-based parcellation map in Franz-Xaver Neubert et al.^53^. All the remaining masks were retrieved from the Harvard-Oxford Atlas distributed with FSL. To create the dorsal striatum masks, we pooled together in one single mask the caudate and putamen masks from the Harvard-Oxford Atlas. For the insula, amygdala, accumbens and dorsal striatum, we decided to consider right and left homologous structures separately as we have previously described some degree of left lateralization of the effects of intranasal OT on rCBF in men^39, 40^. We used a full factorial linear mixed model, including diagnosis, treatment and diagnosis x treatment as fixed effects, participants as a random effect and global grey-matter CBF as a nuisance variable. All analyses were implemented in SPSS24(http://www-01.ibm.com/software/uk/analytics/spss/). Results are reported at a level of significance α = 0.05. For the ROI analyses, we contained the false-discovery rate for the number of ROIs tested at α=0.05 using the Benjamini-Hochberg procedure^54^.

Finally, we conducted a whole-brain exploratory investigation of treatment, diagnosis and treatment x diagnosis effects on rCBF, using global grey-matter CBF as a covariate (see Supplementary Material for details). We used cluster-level inference at α = 0.05 using family-wise error (FWE) correction for multiple comparisons and a cluster-forming threshold of P=0.005 (uncorrected). These statistical thresholds had been determined *a priori* based on our own work investigating the effects of intranasal OT on rCBF in humans^40, 42^ and are standardly applied in ASL studies measuring rCBF^55–60^.

### Associations between clinical measures and rCBF in women with BN/BED and healthy controls

To investigate if changes in rCBF in women with BD/BED were related to the severity of clinical symptomatology, we correlated the mean rCBF in each of the four anatomical regions-of-interest where we found significant differences between the BN/BED and healthy groups with clinical measures. We focused our analyses on mean rCBF values extracted from anatomical regions-of-interest to avoid potential issues of selection bias if we based our analyses on rCBF extracted from clusters showing significant diagnostic group differences in the whole brain analyses. As a measure of eating symptom severity we used the global EDEQ scores. Since patients with BN/BED score highly on scales measuring anxiety, stress and depression, and scores on these scales were highly correlated with each other, we used within-group principal component analysis (PCA) on these three measures to obtain a single score reflecting affective/stress symptom severity. The first principal component accounted for 70.81% and 83.47% of the total variance in the patients and healthy controls, respectively and was used in our analyses to examine the specificity of the association of rCBF with eating symptom severity.

We estimated partial Pearson correlation coefficients (with bootstrapping – 1000 samples) between mean rCBF and global EDEQ or the first component of the stress, anxiety and depression scores in each region-of-interest, adjusting for global CBF and BMI. To examine the specificity of the association between mean rCBF and eating symptomatology, we re-estimated the partial correlations with global EDEQ after including scores on the first principal component from the PCA on the stress, anxiety and depression measures. We estimated these correlations separately in patients and healthy controls because group differences in mean scores on these measures might result in illusory correlations if the two samples were pooled together^62^. For completeness, we then compared these correlations between the patient and healthy control groups using the Fisher r-to-z transformation.

### Post-hoc analyses

#### BMI

Since BMI influences perfusion in the brain^63^ and patients presented high variability in BMI, we repeated all our analyses including BMI as a nuisance variable to account for BMI related variability in rCBF.

#### Hormonal contraception

Hormonal contraception can reduce intranasal OT-induced effects on brain physiology, at least in response to social stimuli^64^. Since 37.5% of our women were under hormonal contraception, we repeated all analyses including hormonal contraception as a nuisance variable.

#### Diagnostic category, psychiatric comorbidities and current pharmacological treatment

To disentangle whether the main effect of diagnosis was driven specifically by BED status, current pharmacological treatment or presence of other psychiatric conditions, we repeated our analyses for the main effect of diagnosis either excluding the 5 BED patients or including diagnostic category and comorbidities/current treatment as nuisance variables.

#### Grey-matter volume

Variations in grey-matter volume (GMV) can be associated with local variations in brain tissue metabolic demand, therefore influencing rCBF^65, 66^. For this reason, we explored if the main effects of diagnosis were related to differences in GMV. For each participant, we used GM volume probability maps obtained after segmentation of the T1-weighted structural image and the *fslmeants* command to estimate mean GMV values in ROIs based on the clusters where BN/BED patients showed significant increases in rCBF compared to controls in the whole-brain analyses (details provided in Supplementary Material). We then repeated the whole-brain analysis testing for the main effects of diagnosis on rCBF including GMV as a covariate. For completeness, we also ran an exploratory whole-brain analysis testing for the main effect of diagnosis on GMV (further details in Supplementary Material). Our main motivation was to investigate if clusters showing significant effects of diagnosis in rCBF mapped onto brain areas showing significant GMV abnormalities, giving us insight in the interpretation of rCBF abnormalities in BN/BED.

## RESULTS

### Sample characteristics

Women with BN/BED and healthy controls did not differ in age, height, BMI, educational level, or hormonal status (see table 1). Women with BN/BED had significantly higher weight, anxiety, depression and stress scores, and EDEQ global and subscale scores compared to healthy women. Women with BN/BED had an average duration of eating disorder of 10.15 years, and the average age of onset was 15.90 years old.

**Table 1.**
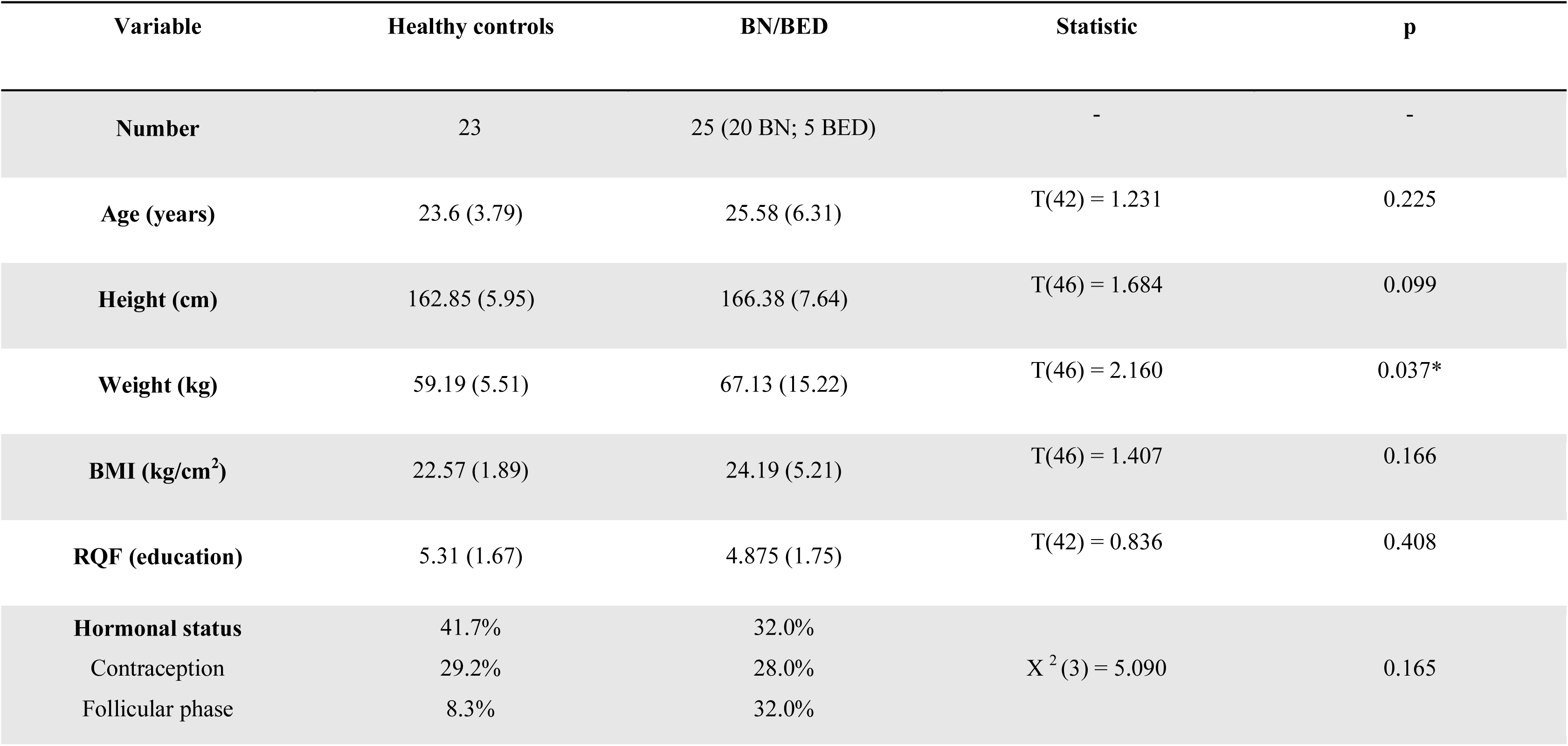
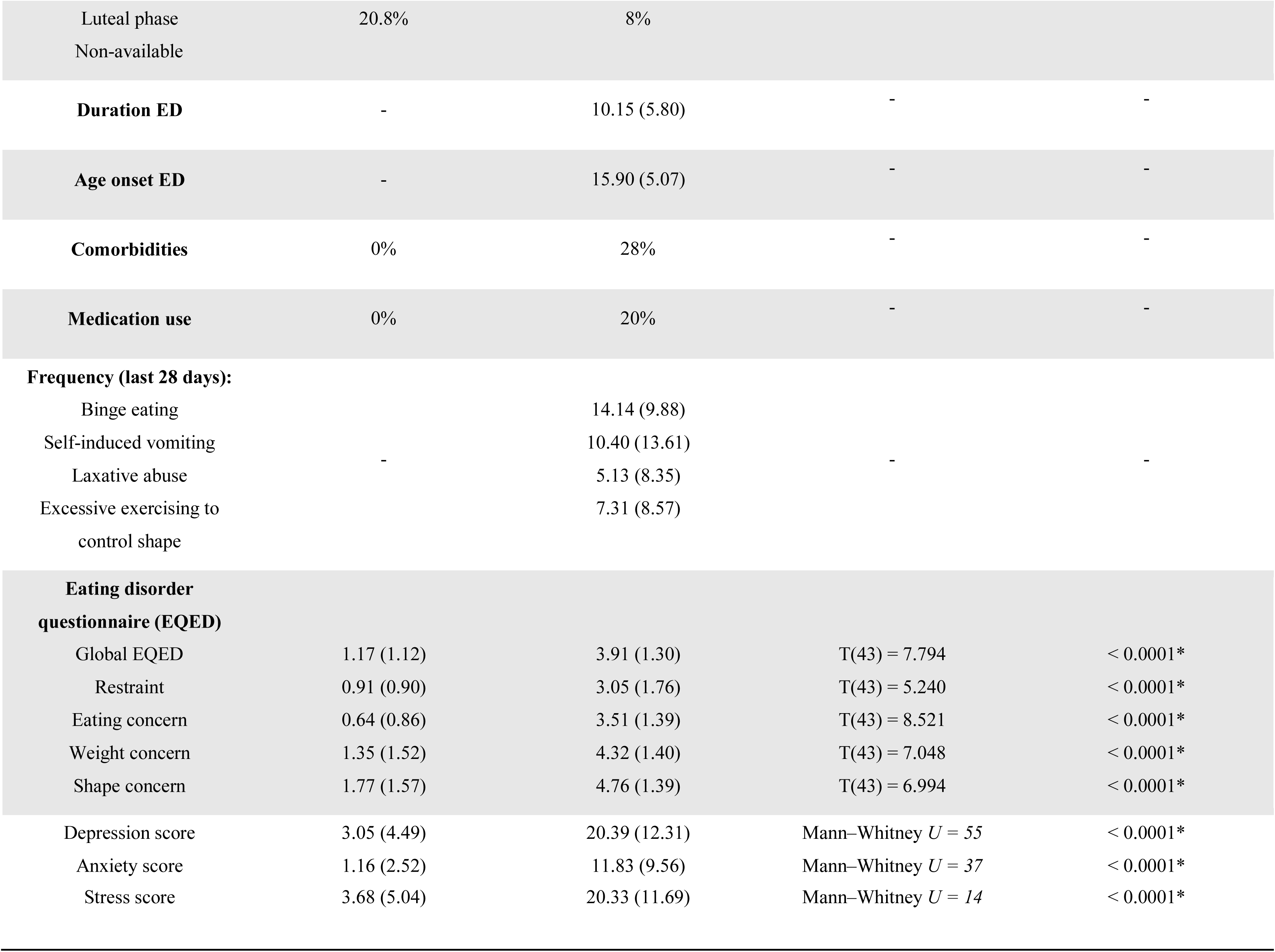
Sociodemographic characteristics. This table shows the sociodemographic characteristics of our cohort of healthy and BN/BED women. Data are presented as mean (standard deviation). The fourth and fifth columns summarize the statistics corresponding to groups comparison for each variable, when applicable. Statistical significance was set to *p* < 0.05 and is highlighted with the symbol *.

### Diagnosis, treatment and diagnosis x treatment effects on resting global CBF

There were no effects of diagnosis, treatment or diagnosis x treatment on global grey-matter CBF (Fig. S2).

### Diagnosis, treatment and diagnosis x treatment effects on resting rCBF

#### ROI analyses

Women with BN/BED showed significant rCBF increases in the medial prefrontal cortex (PFC) and orbitofrontal cortex (OFC) ROIs compared to healthy controls (Table S1). These differences did not survive correction for the number of total ROIs tested. Accounting for BMI, women with BN/BED showed significant rCBF increases in the medial PFC, OFC, right insula and anterior cingulate gyrus (ACG) ROIs (Table 2 and Fig. 1) that remained significant after correcting for multiple testing.

**Fig. 1.**
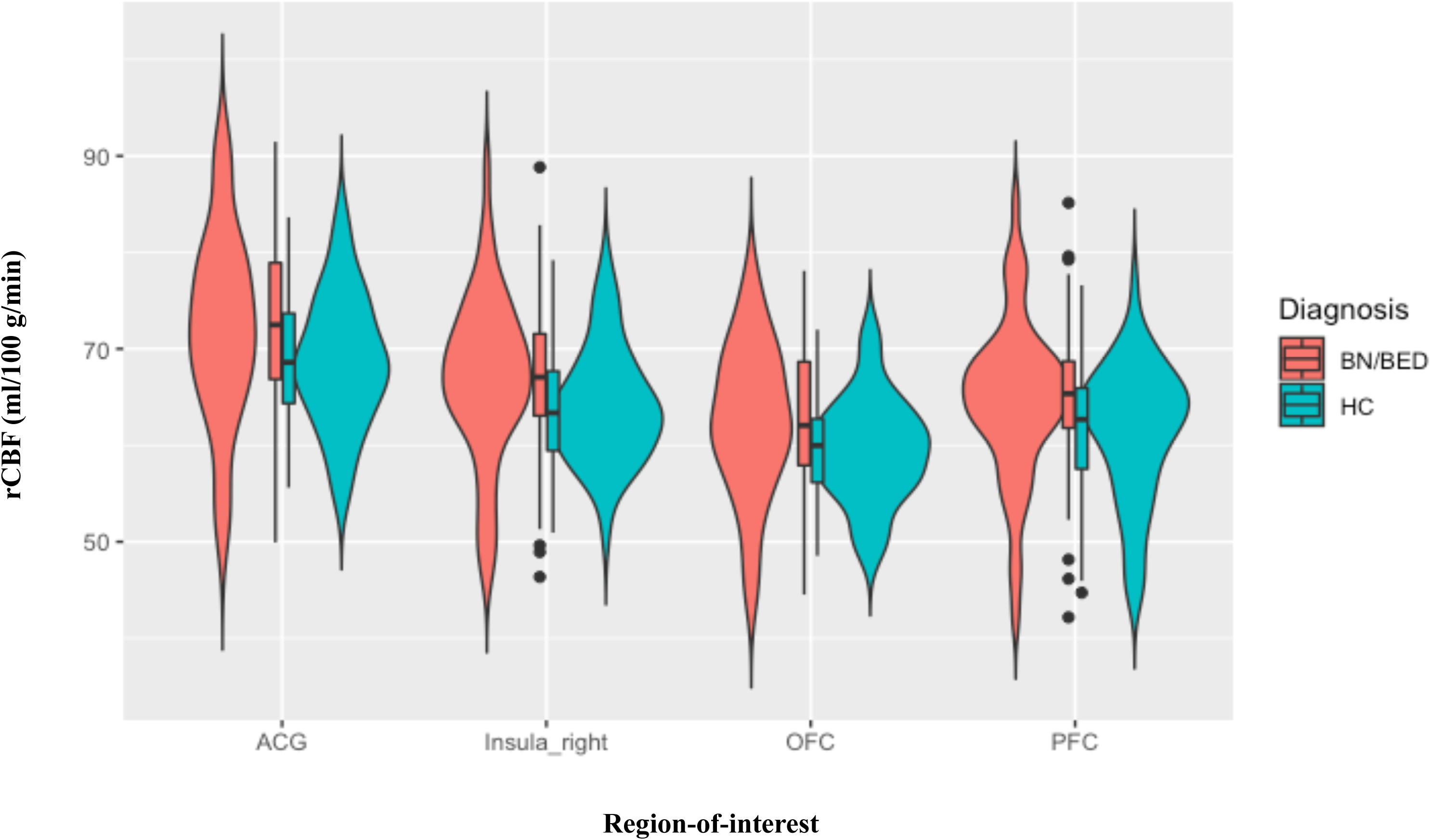
Increases in mean resting regional cerebral blood flow (rCBF) in the anterior cingulate, right insula, orbitofrontal and medial prefrontal cortices in BN/BED women compared to healthy women (hypothesis-driven analysis). These graphs illustrate the changes in mean rCBF in BN/BED patients compared to healthy controls for the regions-of-interest where we identified a significant main effect of diagnosis. For all of these four regions-of-interest, BN/BED women presented higher mean rCBF than healthy women; ACG – anterior cingulate gyrus; OFC – orbitofrontal cortex; PFC – medial prefrontal cortex; Box plots and violin plots depicting mean rCBF (marginal means) on each region-of-interest for each diagnosis/treatment groups; middle horizontal lines represent the median; boxes indicate the 25^th^ and 75^th^ percentiles.

**Table 2.**
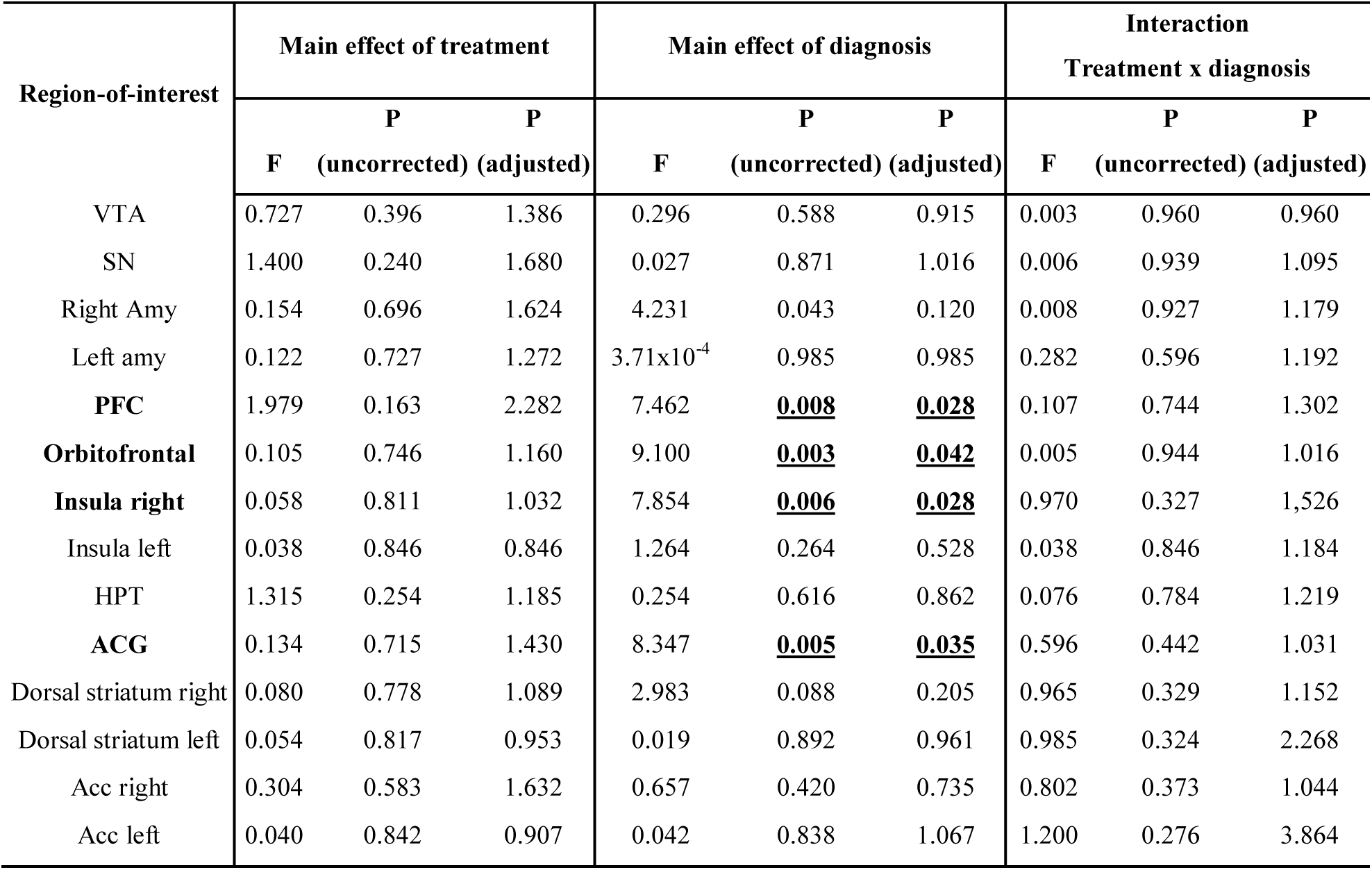
Effects of diagnosis, treatment and diagnosis x treatment on resting regional cerebral blood flow (rCBF) within neural circuits relevant for BN/BED (hypothesis-driven analysis). This table shows the results of a hypothesis-driven investigation of the effects of diagnosis, treatment and diagnosis x treatment on rCBF within 14 anatomical regions-of-interest suggested to be involved in BN/BED. We tested these effects in a liner mixed model, controlling for global grey-matter cerebral blood flow and BMI. Statistical significance was set to *p* < 0.05, after correction for multiple testing with the Benjamini-Hochberg procedure; VTA – Ventral tegmental area; SN – Substantia nigra; Amy – Amygdala; PFC – Prefrontal Cortext; HPT – Hypothalamus; ACG – Anterior Cingulate Gyrus; Acc – Accumbens).

#### Whole brain analysis

Exploratory whole-brain analysis revealed rCBF increases in BN/BED patients in 3 clusters (Fig. S3 and Table S2). Two clusters spanned the inferior/middle temporal gyrus bilaterally (with a bigger extent at the right lobe) and one cluster the PFC, medial OFC and ACG bilaterally (extending more into the left hemisphere). Accounting for BMI extended the right inferior/middle temporal gyrus cluster to include the right posterior insula (Fig. 2 and Table S3). We did not observe any treatment or treatment x diagnosis effects.

**Fig. 2.**
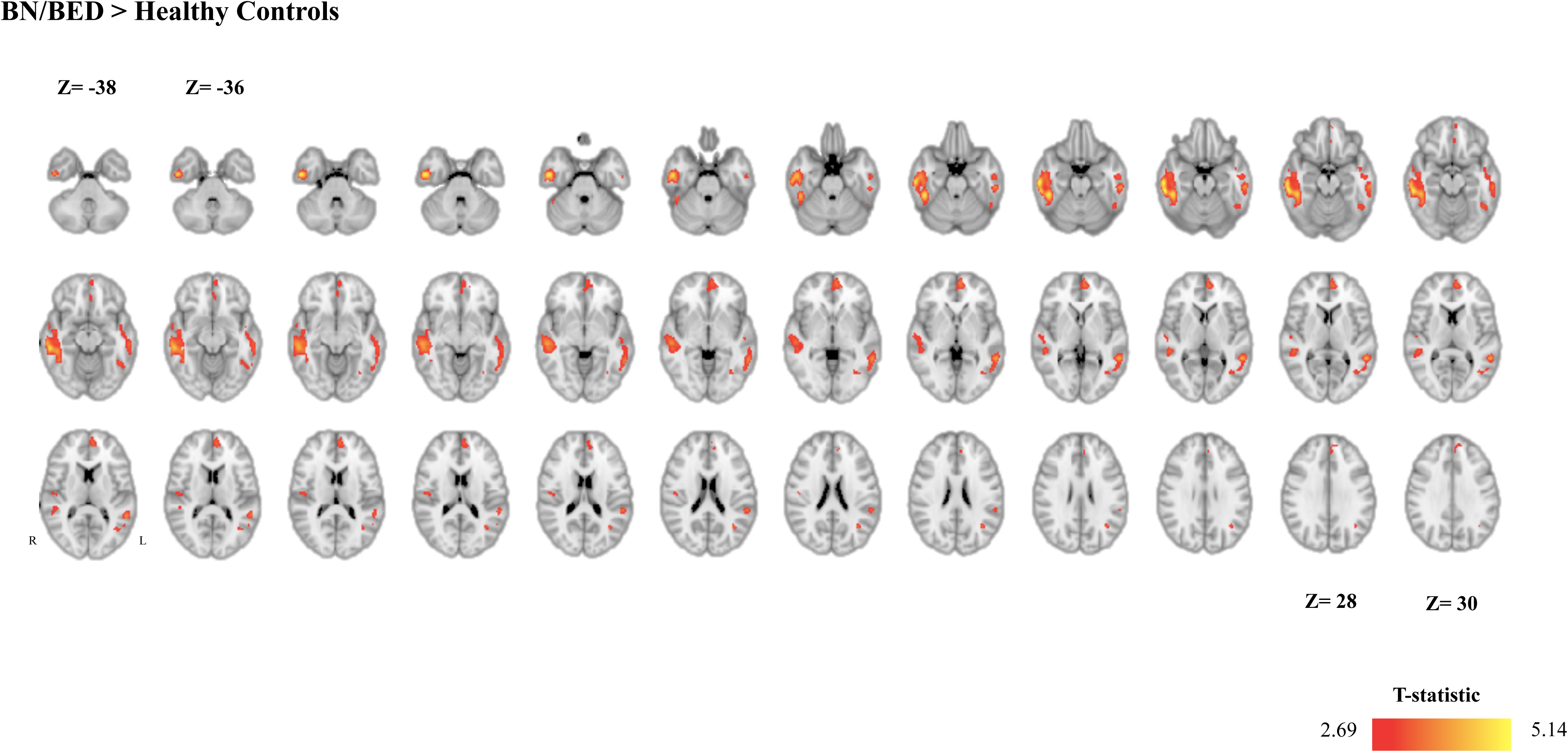
Increases in resting regional cerebral blood flow (rCBF) in the brain of BN/BED patients (whole-brain analysis). This figure shows the results of a directed T-contrast analysis at the whole-brain level where we tested for increases (BN/BED > Controls) or decreases (Controls > BN/BED) in rCBF in patients compared to controls, accounting for global grey-matter cerebral blood flow and BMI. Whole-brain cluster-level inference was applied at α = 0.05 using familywise error (FWE) correction for multiple comparisons and a cluster-forming threshold of *p* = 0.005 (uncorrected). Images are shown as T-statistic in radiological convention. We did not find any significant cluster BN/BED patients presented lower rCBF than healthy controls.

### Associations between clinical symptoms and rCBF in women with BN/BED and healthy controls

We observed significant positive partial correlations (adjusting for BMI and global CBF) between mean rCBF extracted from the medial PFC, OFC, ACG and right insula ROIs and global EDEQ scores, in patients but not in controls (Fig. 3). Partial correlations between mean rCBF in these ROIs and global EDEQ were still significant after additionally accounting for stress, anxiety and depression (Table S4), except for the medial PFC ROI for which the correlation was at trend-level (p=0.051). However, direct comparisons of these correlations between the two groups yielded no statistically significant differences (Fig. 3 and Table S4).

**Fig. 3.**
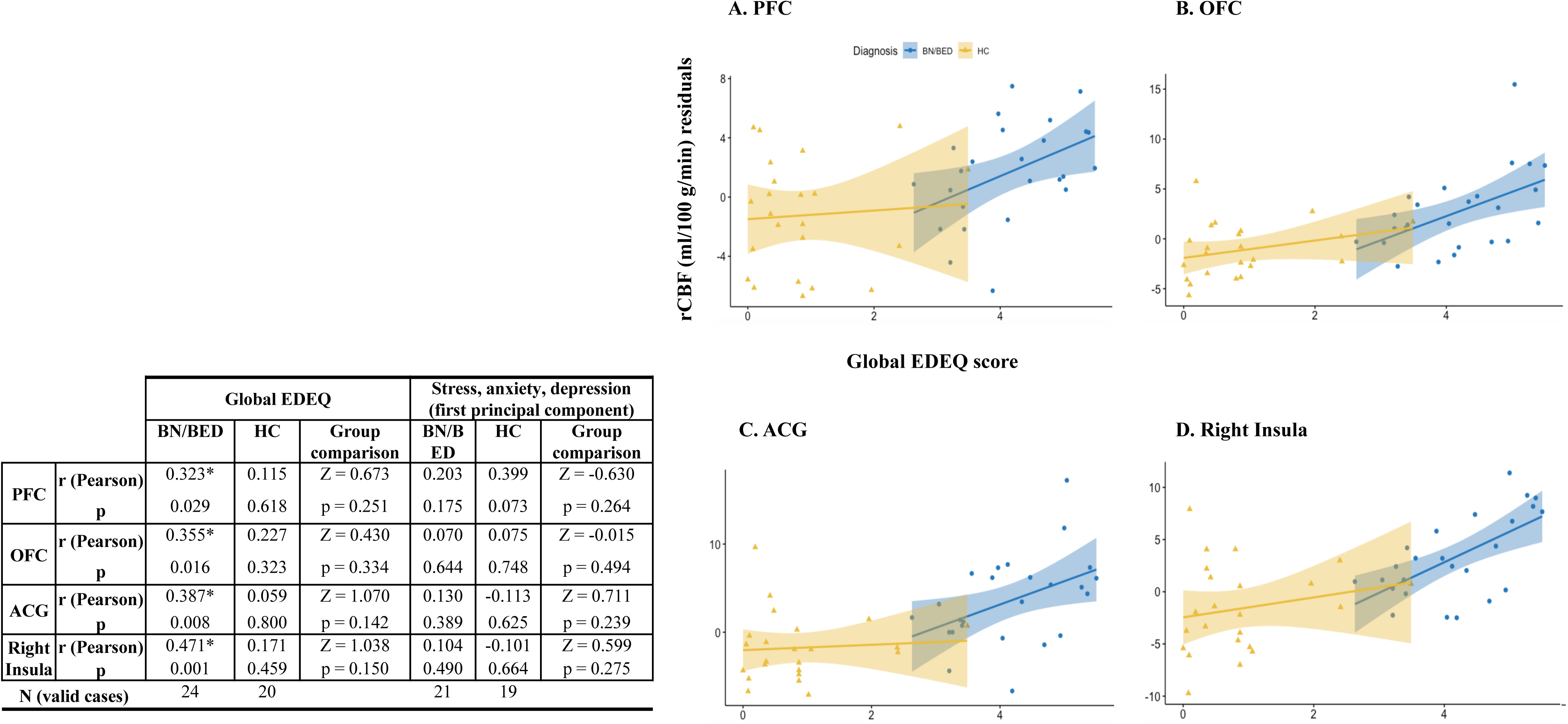
Increases in resting regional cerebral blood flow (rCBF) in the brain of BN/BED patients correlate positively with eating disorder symptom severity. *Left panel* - Partial Pearson correlations between mean rCBF in each of the four anatomical regions-of-interest where we found significant differences between the BN/BED and healthy groups and clinical symptomatology. We used the global EDEQ scores as a measure of eating disorder symptom severity and the first principal component of the anxiety, stress and depression scores. Partial Pearson correlations were calculated with bootstrapping (1000 samples) adjusting for global CBF and BMI, separately for controls and patients. In the last columns of each sub-section, we present the result of the statistical comparison of the correlations between the two groups, as assessed by Fisher r-to-z transformation. *Right panel* – We present scatter plots depicting the relationship between mean rCBF (marginal scores after regressing out the effects of global CBF and BMI in the y-axis) and global EDEQ scores (in the x – axis) in each region-of-interest. Statistical significance was set to *p* < 0.05 and is highlighted with the symbol *. PFC – Medial Prefrontal cortex; OFC – Orbitofrontal cortex; ACG – Anterior Cingulate Gyrus.

### Post-hoc analyses

#### Hormonal contraception

Accounting for use of hormonal contraception produced no change in our findings (Tables S5, Fig. S4).

#### Diagnostic category, psychiatric comorbidities and current pharmacological treatment

Excluding the BED patients (Fig. S5) or accounting for diagnostic category (Fig. S6) as a covariate in our model produced did not substantially change our results. We also did not find any substantial changes in our results when we accounted for comorbidities/current drug treatment (Fig. S7).

### Is regional grey matter volume (GMV) associated with the effect of diagnosis in rCBF?

The main effect of diagnosis in the whole brain analysis remained virtually unaltered after accounting for GMV values in the ROIs based on the clusters where BN/BED patients showed significant increases in rCBF compared to controls in the whole-brain analyses (Fig. S8). Furthermore, women with BN/BED showed lower GMV than controls in two clusters, one spanning the postcentral and precentral gyri and another the inferior/middle temporal gyrus, both in the right hemisphere (Fig. 4). Interestingly, the right temporal gyrus cluster partially overlapped with the cluster showing rCBF increases in women with BD/BED compared to controls (Fig. S9). In the right temporal gyrus, rCBF and GMV correlated negatively in patients but not in controls (Fig. S10) (BN/BED: r=-0.496, p=0.012; healthy controls: r=-0.024, p=0.913; Groups comparison: z=-1.66, p=0.048). We did not identify any area in the brain where patients presented higher GMV than controls.

## DISCUSSION

Using ASL, a highly reproducible and widely available quantitative MRI technique, we demonstrated, for the first time, resting hyperperfusion abnormalities in women with BN/BED, compared to healthy controls, in key neural circuits implicated in BN/BED psychopathology. Hyperperfusion abnormalities in the PFC, medial OFC, ACG and the right posterior insula were positively and specifically associated with symptom severity as reflected on the global EDEQ scale in women with BN/BED. However, a single dose of 40 IU intranasal oxytocin did not restore or attenuate these perfusion abnormalities at 18-26 minutes post-dosing. Our findings enhance our understanding of resting brain abnormalities in BN/BED and identify resting rCBF as a non-invasive potential biomarker for disease-related changes and treatment monitoring. We discuss each of our main findings below in turn.

### Abnormalities in resting brain perfusion in women with BN/BED compared to healthy controls

Our hypothesis-driven analyses showed that women with BN/BED, compared to healthy controls, presented with increased resting rCBF in the medial PFC, medial OFC, ACG and the right posterior insula. Exploratory whole-brain analysis showed further increases in the inferior/middle/superior temporal cortices in women with BN/BED. Sensitivity analyses confirmed that these findings were not driven by the inclusion of BED patients and did not reflect current medication or other psychiatric comorbidities. In women with BN/BED, estimates of rCBF in three key areas where we found women with BN/BED and healthy women to differ in rCBF (medial PFC, ACG and right insula) correlated positively with global scores on the EDEQ, even after adjusting for anxiety, depression and stress, supporting a specific association between rCBF abnormalities in these regions and the underlying eating disorder psychopathology. Given the high reliability and ease of acquisition, our findings suggest that measuring resting rCBF at BN/BED patients with ASL MRI can offer a non-invasive *in-vivo* biomarker of potential value for understanding disease-related functional changes and to examine treatment effects, posing minimal demand and no risk on patients.

We are aware of only three studies examining rCBF in BN/BED, using SPECT and small sample sizes (7<n<14). One study found increased rCBF in the right temporal lobe in response to own body images when a pooled sample of patients with DSM-IV BN – purging type and anorexia nervosa – restrictive type were compared to a pooled sample of healthy controls, patients with DSM-IV BN – non-purging type and patients with anorexia nervosa– purging type^17^. Another study reported increased rCBF in the left temporal lobe and inferior frontal region bilaterally in patients with DSM-III BN before eating^16^. Finally, the only study examining patients with BED reported that obese BED patients, compared to non-obese BED patients and healthy controls, presented increased rCBF responses in the frontal and prefrontal cortices in response to food visual stimuli^18^. These studies did not report on rCBF changes at rest, which precludes direct comparisons between these studies and ours. Nevertheless, we note that the changes in rCBF reported in these studies are in the same direction and largely overlap with regions where we found resting hyperperfusion during BN/BED in our study. Therefore, it is possible that hyperperfusion in these areas in women with BN/BED marks a disease-relevant state which can be captured at rest and without the need of a specific experimental manipulation.

The mechanisms behind these increases in resting rCBF in BN/BED remain unknown. Increases in resting rCBF are likely to reflect basal regional hypermetabolism, potentially associated with increases in resting neural activity^15^. We note however that the few small-scale studies evaluating resting brain glucose metabolism using PET during BN have generally reported decreases in global and regional glucose metabolism, including parietal and anterior frontal hypometabolism^67–71^. The only exception is one study that found increased glucose metabolism in the temporal lobes of patients with BN, which matches our findings of increased perfusion in these areas^69^. BN patients have been reported to present higher GMV in some areas such as the medial orbitofrontal cortex^8^, which could account for the increases in rCBF we report herein; however, we found that this is unlikely to be the case in our sample. Exploratory whole-brain analysis showed decreased GMV in the right temporal lobe in BN/BED compared to controls. This cluster partially overlapped with one cluster showing increased rCBF in patients. GMV was negatively correlated with rCBF in this area in patients, but not in controls. We speculate that two possible mechanisms could account for this paradoxical relationship between rCBF and GMV in the temporal lobe of patients with BN/BED. One possibility is that the increases in rCBF we observed in the right temporal lobe might reflect a mechanism of local functional plasticity in response to GMV loss in this region - which ultimately may help to maintain temporal lobe function in patients with BN/BED^72, 73^. Alternatively, it is also conceivable that enduring neural hyperactivity, accompanied by increases in rCBF, could drive GMV loss due to excitotoxicity^74^. Future longitudinal studies examining how changes in GMV and rCBF in these brain regions develop may elucidate these hypotheses.

Since our data were acquired at rest, we can only speculate regarding the specific contribution of the perfusion and structural abnormalities observed in this study to core behavioural manifestations in BN/BED. We note that BN/BED patients have been reported to present with enhanced sensitivity to the anticipation of food reward/hedonic value^5^ which matches the increases in rCBF we observed in the medial OFC (a key area involved in reward/outcome value processing^75^, including the hedonic value of food^76^). BN, in particular, and to some extent BED, have also been associated with continuous feelings of monitoring of binge-activating stimuli in the environment and body-shape/weight self-judgement, which match our findings of increased rCBF in the medial PFC and ACC (two key areas involved in self-monitoring and control regulation^77^) in patients. Evidence from lesion studies have supported a direct implication of the temporal lobe (predominantly the right lobe) in eating disorders^78^. While the exact mechanisms by which temporal lobe dysfunction may contribute to eating dysregulation remain elusive, some studies have implicated the superior/middle temporal gyri in the brain responses to palatable food^79^ and in inhibitory control, including cognitive control of appetite^80^ – two core elements of binge-eating. Importantly, previous studies have reported attenuated recruitment of these areas during response inhibition in both BN^81, 82^ and BED^83^.

In contrast to our hypotheses, we could not find any resting rCBF alterations in the ventral or dorsal striatum in patients with BN/BED. This is surprising, given the pivotal role of the striatal circuits in cognitive processes known to be disrupted in BN/BED (such as incentive and habitual behaviours^84^, impulsivity^85^ and self-regulation^86^) and previous neuroimaging studies demonstrating functional and structural abnormalities in these areas (see ^7, 8^ for detailed reviews). It is possible that while disrupted rCBF in the striatum in BN/BED may not manifest at rest, alterations may become evident in condition engaging this region, such as the anticipation or valuation of hedonic stimuli (e.g. food).

### Does intranasal OT restore rCBF abnormalities in women with BN/BED**?**

The lack of treatment or treatment x diagnosis effects does not support a normalizing effect of 40IU intranasal OT on resting rCBF abnormalities in women with BN/BED 18-26 mins post-dosing. A number of reasons might explain the lack of effects of intranasal OT on rCBF in this study. First, it is possible that we may have missed the active time-window for intranasal OT treatment effects in women with BN/BED. We have previously mapped intranasal OT-induced effects on brain perfusion in men and demonstrated that while 40IU intranasal OT can induce changes in resting perfusion as early as 15 minutes post-doing (earlier intervals have not been sampled), the effects do vary as a function of method of administration, dose, and the latency of the sampling interval post-dosing^40^. However, we are still lacking an in-depth pharmacodynamics investigation of OT-induced changes in rCBF in women. Previous studies have shown that the effects of the administration of the same dose of intranasal OT to men and women can result in different effects on behaviour and brain function^87–91^. A recent study has shown that differences in the effects of a range of doses of intranasal OT on brain responses to happy and fearful faces between men and women do not simply reflect differences in dose sensitivity, but rather gender-specific effects^92^. Therefore, while we have informed this current study based on our in-depth characterizations of the pharmacodynamics of intranasal OT in men, it is possible that crucial differences exist in the spatiotemporal pattern of rCBF changes after intranasal OT between genders. Future studies should systematically investigate the potential effects of these factors, including a comparison of acute versus chronic regimens of administration, characterization of rCBF changes over an extended period of time and dose-response studies in women. Second, hormonal contraception has been shown to blunt responses to intranasal OT in women^64^. Although accounting for contraception in our analyses produced virtually no change in our results, it is plausible that OT might have exerted an effect on rCBF in our sample in the absence of hormonal contraception. Finally, while we could not detect any significant treatment effects on rCBF in women with or without BN/BED at rest in this study, we cannot exclude that significant effects of intranasal OT might emerge during targeted experimental challenges, such as exposure to food cues^93^ or stress^94^.

## Limitations

One limitation of our study is that we only included women. Our findings should thus not be extrapolated to men with BN/BED. Secondly, we could only enrol a small number of BED patients, which did not allow us to isolate effects related to diagnostic category. Thirdly, while we have asked subjects to eat 2.5h before our experimental sessions in order to minimize the effects of baseline hunger, we did not standardized the amount or type of food consumed – which may have introduced some noise in our data.

## Conclusions

BN/BED in women is accompanied by increased resting perfusion in key brain areas potentially associated with the underlying eating psychopathology. Intranasal OT did not attenuate or restore these abnormalities, at least for the specific dose and post-dosing interval examined. Future studies examining a more comprehensive range of doses, time-windows and schemes of administration will be needed before we can ascertain the therapeutic value of intranasal OT in improving resting rCBF disturbances in women with BN/BED. Given the high reliability and ease of acquisition, measuring resting rCBF at BN/BED patients with ASL MRI offers a promising non-invasive *in-vivo* biomarker of functional changes in these patients, with potential implications for diagnosis and treatment monitoring.

## ACKNOWLEDGMENTS

The authors wish to thank the study volunteers for their participation.

## ADDITIONAL INFORMATION

### Author contributions

YP and JT designed the study; ML collected the data; DM performed data analysis; SR and FZ contributed to data analysis; DM and YP interpreted the data and drafted the first version of the manuscript; all authors (DM, ML, SR, FZ, JT, YP) provided critical revisions approved the final version of the manuscript to be published.

### Data availability

Data can be provided upon request.

## REFERENCES

1. Smith, K. E. et al. The validity of DSM-5 severity specifiers for anorexia nervosa, bulimia nervosa, and binge-eating disorder. Int J Eat Disord 50, 1109–1113, doi:10.1002/eat.22739 (2017).

2. Alvarenga, M. S. et al. Eating attitudes of anorexia nervosa, bulimia nervosa, binge eating disorder and obesity without eating disorder female patients: differences and similarities. Physiol Behav 131, 99–104, doi:10.1016/j.physbeh.2014.04.032 (2014).

3. Aguera, Z. et al. Short-Term Treatment Outcomes and Dropout Risk in Men and Women with Eating Disorders. European eating disorders review : the journal of the Eating Disorders Association 25, 293–301, doi:10.1002/erv.2519 (2017).

4. Turton, R., Chami, R. & Treasure, J. Emotional Eating, Binge Eating and Animal Models of Binge-Type Eating Disorders. Curr Obes Rep 6, 217–228, doi:10.1007/s13679-017-0265-8 (2017).

5. Schienle, A., Schafer, A., Hermann, A. & Vaitl, D. Binge-eating disorder: reward sensitivity and brain activation to images of food. Biol Psychiatry 65, 654–661, doi:10.1016/j.biopsych.2008.09.028 (2009).

6. Stojek, M. et al. A systematic review of attentional biases in disorders involving binge eating. Appetite 123, 367–389, doi:10.1016/j.appet.2018.01.019 (2018).

7. Kessler, R. M., Hutson, P. H., Herman, B. K. & Potenza, M. N. The neurobiological basis of binge-eating disorder. Neurosci Biobehav Rev 63, 223–238, doi:10.1016/j.neubiorev.2016.01.013 (2016).

8. Donnelly, B. et al. Neuroimaging in bulimia nervosa and binge eating disorder: a systematic review. J Eat Disord 6, 3, doi:10.1186/s40337-018-0187-1 (2018).

9. Domakonda, M. J., He, X., Lee, S., Cyr, M. & Marsh, R. Increased Functional Connectivity Between Ventral Attention and Default Mode Networks in Adolescents With Bulimia Nervosa. J Am Acad Child Adolesc Psychiatry 58, 232–241, doi:10.1016/j.jaac.2018.09.433 (2019).

10. Amianto, F. et al. Intrinsic connectivity networks within cerebellum and beyond in eating disorders. Cerebellum 12, 623–631, doi:10.1007/s12311-013-0471-1 (2013).

11. Wang, L. et al. Altered intrinsic functional brain architecture in female patients with bulimia nervosa. J Psychiatry Neurosci 42, 414–423 (2017).

12. Dunlop, K. et al. Increases in frontostriatal connectivity are associated with response to dorsomedial repetitive transcranial magnetic stimulation in refractory binge/purge behaviors. Neuroimage Clin 8, 611–618, doi:10.1016/j.nicl.2015.06.008 (2015).

13. Lee, S. et al. Resting-state synchrony between anterior cingulate cortex and precuneus relates to body shape concern in anorexia nervosa and bulimia nervosa. Psychiatry Res 221, 43–48, doi:10.1016/j.pscychresns.2013.11.004 (2014).

14. Turner, R. Uses, misuses, new uses and fundamental limitations of magnetic resonance imaging in cognitive science. Philosophical transactions of the Royal Society of London. Series B, Biological sciences 371, doi:10.1098/rstb.2015.0349 (2016).

15. Paulson, O. B., Hasselbalch, S. G., Rostrup, E., Knudsen, G. M. & Pelligrino, D. Cerebral blood flow response to functional activation. J Cereb Blood Flow Metab 30, 2–14, doi:10.1038/jcbfm.2009.188 (2010).

16. Nozoe, S. et al. Comparison of regional cerebral blood flow in patients with eating disorders. Brain Res Bull 36, 251–255 (1995).

17. Beato-Fernandez, L., Rodriguez-Cano, T. & Garcia-Vilches, I. Psychopathological alterations and neuroimaging findings with discriminant value in eating behavior disorders. Actas Esp Psiquiatr 39, 203–210 (2011).

18. Karhunen, L. J. et al. Regional cerebral blood flow during exposure to food in obese binge eating women. Psychiatry Res 99, 29–42 (2000).

19. Bateman, T. M. Advantages and disadvantages of PET and SPECT in a busy clinical practice. J Nucl Cardiol 19, S3–S11, doi:10.1007/s12350-011-9490-9 (2012).

20. Hodkinson, D. J. et al. Quantifying the test-retest reliability of cerebral blood flow measurements in a clinical model of on-going post-surgical pain: A study using pseudo-continuous arterial spin labelling. Neuroimage Clin 3, 301–310, doi:10.1016/j.nicl.2013.09.004 (2013).

21. Wolf, R. L. & Detre, J. A. Clinical neuroimaging using arterial spin-labeled perfusion magnetic resonance Imaging. Neurotherapeutics 4, 346–359, doi:DOI 10.1016/j.nurt.2007.04.005 (2007).

22. Peterson, B. S. et al. Hyperperfusion of Frontal White and Subcortical Gray Matter in Autism Spectrum Disorder. Biol Psychiatry, doi:10.1016/j.biopsych.2018.11.026 (2018).

23. Ota, M. et al. Pseudo-continuous arterial spin labeling MRI study of schizophrenic patients. Schizophr Res 154, 113–118, doi:10.1016/j.schres.2014.01.035 (2014).

24. Colloby, S. J. et al. Regional cerebral blood flow in late-life depression: arterial spin labelling magnetic resonance study. Br J Psychiatry 200, 150–155, doi:10.1192/bjp.bp.111.092387 (2012).

25. Wang, D. J. J., Chen, Y. F., Fernandez-Seara, M. A. & Detre, J. A. Potentials and Challenges for Arterial Spin Labeling in Pharmacological Magnetic Resonance Imaging. Journal of Pharmacology and Experimental Therapeutics 337, 359–366, doi:10.1124/jpet.110.172577 (2011).

26. Stewart, S. B., Koller, J. M., Campbell, M. C. & Black, K. J. Arterial spin labeling versus BOLD in direct challenge and drug-task interaction pharmacological fMRI. Peerj 2, doi:ARTN e687 10.7717/peerj.687 (2014).

27. Plessow, F., Eddy, K. T. & Lawson, E. A. The Neuropeptide Hormone Oxytocin in Eating Disorders. Curr Psychiatry Rep 20, 91, doi:10.1007/s11920-018-0957-0 (2018).

28. Giel, K., Zipfel, S. & Hallschmid, M. Oxytocin and Eating Disorders: A Narrative Review on Emerging Findings and Perspectives. Current Neuropharmacology 16, 1111–1121, doi:10.2174/1570159x15666171128143158 (2018).

29. Romano, A., Tempesta, B., Micioni Di Bonaventura, M. V. & Gaetani, S. From Autism to Eating Disorders and More: The Role of Oxytocin in Neuropsychiatric Disorders. Front Neurosci 9, 497, doi:10.3389/fnins.2015.00497 (2015).

30. Leslie, M., Silva, P., Paloyelis, Y., Blevins, J. & Treasure, J. A Systematic Review and Quantitative Meta-Analysis of Oxytocin’s Effects on Feeding. Journal of neuroendocrinology, e12584 (2018).

31. Thienel, M. et al. Oxytocin’s inhibitory effect on food intake is stronger in obese than normal-weight men. International Journal of Obesity 40, 1707–1714, doi:10.1038/ijo.2016.149 (2016).

32. Ott, V. et al. Oxytocin Reduces Reward-Driven Food Intake in Humans. Diabetes 62, 3418–3425, doi:10.2337/db13-0663 (2013).

33. Ott, V. et al. Oxytocin reduces reward-driven food intake in humans. Diabetes 62, 3418–3425, doi:10.2337/db13-0663 (2013).

34. Kim, Y. R., Eom, J. S., Yang, J. W., Kang, J. & Treasure, J. The Impact of Oxytocin on Food Intake and Emotion Recognition in Patients with Eating Disorders: A Double Blind Single Dose Within-Subject Cross-Over Design. PLoS One 10, e0137514, doi:10.1371/journal.pone.0137514 (2015).

35. Kim, Y. R., Eom, J. S., Leppanen, J., Leslie, M. & Treasure, J. Effects of intranasal oxytocin on the attentional bias to emotional stimuli in patients with bulimia nervosa. Psychoneuroendocrinology 91, 75–78, doi:10.1016/j.psyneuen.2018.02.029 (2018).

36. Micali, N., Crous-Bou, M., Treasure, J. & Lawson, E. A. Association Between Oxytocin Receptor Genotype, Maternal Care, and Eating Disorder Behaviours in a Community Sample of Women. European eating disorders review : the journal of the Eating Disorders Association 25, 19–25, doi:10.1002/erv.2486 (2017).

37. Kim, Y. R., Kim, J. H., Kim, C. H., Shin, J. G. & Treasure, J. Association between the oxytocin receptor gene polymorphism (rs53576) and bulimia nervosa. European eating disorders review : the journal of the Eating Disorders Association 23, 171–178, doi:10.1002/erv.2354 (2015).

38. Davis, C., Patte, K., Zai, C. & Kennedy, J. L. Polymorphisms of the oxytocin receptor gene and overeating: the intermediary role of endophenotypic risk factors. Nutrition & Diabetes 7, doi:10.1038/nutd.2017.24 (2017).

39. Paloyelis, Y. et al. A Spatiotemporal Profile of In Vivo Cerebral Blood Flow Changes Following Intranasal Oxytocin in Humans. Biological Psychiatry 79, 693–705, doi:10.1016/j.biopsych.2014.10.005 (2016).

40. Martins D, M. N., Zelaya F, Vasilakopoulou S, Loveridge J, Oates A, Maltezos S, Mehta M, Howard M, McAlonan G, Murphy D, Williams S, Fotopoulou A, Schuschnig U, Paloyelis Y. Do direct nose-to-brain pathways underlie intranasal oxytocin-induced changes in regional cerebral blood flow in humans? bioRxiv 563056, doi:https://doi.org/10.1101/563056.

41. Davies, C. et al. Oxytocin modulates hippocampal perfusion in people at clinical high risk for psychosis. Neuropsychopharmacology, doi:10.1038/s41386-018-0311-6 (2019).

42. Paloyelis, Y. et al. A Spatiotemporal Profile of In Vivo Cerebral Blood Flow Changes Following Intranasal Oxytocin in Humans. Biol Psychiatry 79, 693–705, doi:10.1016/j.biopsych.2014.10.005 (2016).

43. Fairburn, C. G. & Beglin, S. J. Assessment of Eating Disorders - Interview or Self-Report Questionnaire. International Journal of Eating Disorders 16, 363–370 (1994).

44. Lovibond, P. F. & Lovibond, S. H. The Structure of Negative Emotional States - Comparison of the Depression Anxiety Stress Scales (Dass) with the Beck Depression and Anxiety Inventories. Behaviour Research and Therapy 33, 335–343, doi:Doi 10.1016/0005-7967(94)00075-U (1995).

45. Chan, C. W. & Leung, S. F. Validation of the Eating Disorder Examination Questionnaire: an online version. J Hum Nutr Diet 28, 659–665, doi:10.1111/jhn.12275 (2015).

46. Shea, T. L., Tennant, A. & Pallant, J. F. Rasch model analysis of the Depression, Anxiety and Stress Scales (DASS). BMC Psychiatry 9, 21, doi:10.1186/1471-244X-9-21 (2009).

47. Fafrowicz, M. et al. Beyond the Low Frequency Fluctuations: Morning and Evening Differences in Human Brain. Front Hum Neurosci 13, 288, doi:10.3389/fnhum.2019.00288 (2019).

48. Kagerbauer, S. M. et al. Absence of a diurnal rhythm of oxytocin and arginine-vasopressin in human cerebrospinal fluid, blood and saliva. Neuropeptides 78, 101977, doi:10.1016/j.npep.2019.101977 (2019).

49. Thedens, D. R., Irarrazaval, P., Sachs, T. S., Meyer, C. H. & Nishimura, D. G. Fast magnetic resonance coronary angiography with a three-dimensional stack of spirals trajectory. Magn Reson Med 41, 1170–1179 (1999).

50. Alsop, D. C. et al. Recommended implementation of arterial spin-labeled perfusion MRI for clinical applications: A consensus of the ISMRM perfusion study group and the European consortium for ASL in dementia. Magn Reson Med 73, 102–116, doi:10.1002/mrm.25197 (2015).

51. Mato Abad, V., Garcia-Polo, P., O’Daly, O., Hernandez-Tamames, J. A. & Zelaya, F. ASAP (Automatic Software for ASL Processing): A toolbox for processing Arterial Spin Labeling images. Magn Reson Imaging 34, 334–344, doi:10.1016/j.mri.2015.11.002 (2016).

52. Pauli, W. M., Nili, A. N. & Tyszka, J. M. Data Descriptor: A high-resolution probabilistic in vivo atlas of human subcortical brain nuclei. Sci Data 5, doi:ARTN 180063 10.1038/sdata.2018.63 (2018).

53. Neubert, F. X., Mars, R. B., Sallet, J. & Rushworth, M. F. S. Connectivity reveals relationship of brain areas for reward-guided learning and decision making in human and monkey frontal cortex. Proceedings of the National Academy of Sciences of the United States of America 112, E2695–E2704, doi:10.1073/pnas.1410767112 (2015).

54. Feser, W. J., Fingerlin, T. E., Strand, M. J. & Glueck, D. H. Calculating Average Power for the Benjamini-Hochberg Procedure. J Stat Theory Appl 8, 325–352 (2009).

55. Mutsaerts, H. et al. Cerebral perfusion changes in presymptomatic genetic frontotemporal dementia: a GENFI study. Brain : a journal of neurology 142, 1108–1120, doi:10.1093/brain/awz039 (2019).

56. Takeuchi, H. et al. Cerebral blood flow during rest associates with general intelligence and creativity. PLoS One 6, e25532, doi:10.1371/journal.pone.0025532 (2011).

57. Joe, A. Y. et al. Response-dependent differences in regional cerebral blood flow changes with citalopram in treatment of major depression. J Nucl Med 47, 1319–1325 (2006).

58. Thomas, B. P. et al. Life-long aerobic exercise preserved baseline cerebral blood flow but reduced vascular reactivity to CO2. J Magn Reson Imaging 38, 1177–1183, doi:10.1002/jmri.24090 (2013).

59. Loggia, M. L. et al. Default mode network connectivity encodes clinical pain: an arterial spin labeling study. Pain 154, 24–33, doi:10.1016/j.pain.2012.07.029 (2013).

60. Nwokolo, M. et al. Hypoglycemic thalamic activation in type 1 diabetes is associated with preserved symptoms despite reduced epinephrine. J Cereb Blood Flow Metab, 271678X19842680, doi:10.1177/0271678X19842680 (2019).

61. Elderkin-Thompson, V. et al. Neuropsychological deficits among patients with late-onset minor and major depression. Am J Geriat Psychiat 10, 60–60 (2002).

62. Paloyelis, Y., Stahl, D. R. & Mehta, M. Are steeper discounting rates in attention-deficit/hyperactivity disorder specifically associated with hyperactivity-impulsivity symptoms or is this a statistical artifact? Biol Psychiatry 68, e15–16, doi:10.1016/j.biopsych.2010.02.025 (2010).

63. Clement, P. et al. Variability of physiological brain perfusion in healthy subjects - A systematic review of modifiers. Considerations for multi-center ASL studies. J Cerebr Blood F Met 38, 1418–1437, doi:10.1177/0271678x17702156 (2018).

64. Scheele, D., Plota, J., Stoffel-Wagner, B., Maier, W. & Hurlemann, R. Hormonal contraceptives suppress oxytocin-induced brain reward responses to the partner’s face. Social Cognitive and Affective Neuroscience 11, 767–774, doi:10.1093/scan/nsv157 (2016).

65. Vasic, N. et al. Baseline brain perfusion and brain structure in patients with major depression: a multimodal magnetic resonance imaging study. J Psychiatry Neurosci 40, 412–421 (2015).

66. Steffener, J., Brickman, A. M., Habeck, C. G., Salthouse, T. A. & Stern, Y. Cerebral blood flow and gray matter volume covariance patterns of cognition in aging. Hum Brain Mapp 34, 3267–3279, doi:10.1002/hbm.22142 (2013).

67. V, D., Goldman, S., V, D. M. & Lotstra, F. Brain glucose metabolism in eating disorders assessed by positron emission tomography. International Journal of Eating Disorders 25, 29–37, doi:Doi 10.1002/(Sici)1098-108x(199901)25:1<29::Aid-Eat4>3.0.Co;2-# (1999).

68. Delvenne, V., Goldman, S., Simon, Y., DeMaertelaer, V. & Lotstra, F. Brain hypometabolism of glucose in bulimia nervosa. International Journal of Eating Disorders 21, 313–320 (1997).

69. Andreason, P. J. Regional Cerebral Glucose-Metabolism in Bulimia-Nervosa (the American Journal of Psychiat, Vol 149, Pg 1509, 1992). American Journal of Psychiatry 150, 174–174 (1993).

70. Krieg, J. C., Holthoff, V., Schreiber, W., Pirke, K. M. & Herholz, K. Glucose-Metabolism in the Caudate Nuclei of Patients with Eating Disorders, Measured by Pet. European Archives of Psychiatry and Clinical Neuroscience 240, 331–333, doi:Doi 10.1007/Bf02279762 (1991).

71. Hagman, J. O. et al. Comparison of Regional Brain Metabolism in Bulimia-Nervosa and Affective-Disorder Assessed with Positron Emission Tomography. Journal of Affective Disorders 19, 153–162, doi:Doi 10.1016/0165-0327(90)90085-M (1990).

72. Wong, T. P. et al. Loss of presynaptic and postsynaptic structures is accompanied by compensatory increase in action potential-dependent synaptic input to layer V neocortical pyramidal neurons in aged rats. Journal of Neuroscience 20, 8596–8606, doi:Doi 10.1523/Jneurosci.20-22-08596.2000 (2000).

73. Hu, F. et al. Use of 3D-ASL and VBM to analyze abnormal changes in brain perfusion and gray areas in nasopharyngeal carcinoma patients undergoing radiotherapy. Biomed Res-India 28, 7879–7885 (2017).

74. Olloquequi, J. et al. Excitotoxicity in the pathogenesis of neurological and psychiatric disorders: Therapeutic implications. Journal of Psychopharmacology 32, 265–275, doi:10.1177/0269881118754680 (2018).

75. Kringelbach, M. L. The human orbitofrontal cortex: linking reward to hedonic experience. Nature reviews. Neuroscience 6, 691–702, doi:10.1038/nrn1747 (2005).

76. Berridge, K. C. ’Liking’ and ‘wanting’ food rewards: brain substrates and roles in eating disorders. Physiol Behav 97, 537–550, doi:10.1016/j.physbeh.2009.02.044 (2009).

77. Heatherton, T. F. Neuroscience of self and self-regulation. Annu Rev Psychol 62, 363–390, doi:10.1146/annurev.psych.121208.131616 (2011).

78. Uher, R. & Treasure, J. Brain lesions and eating disorders. J Neurol Neurosurg Psychiatry 76, 852–857, doi:10.1136/jnnp.2004.048819 (2005).

79. Chen, J., Papies, E. K. & Barsalou, L. W. A core eating network and its modulations underlie diverse eating phenomena. Brain and Cognition 110, 20–42, doi:10.1016/j.bandc.2016.04.004 (2016).

80. Tuulari, J. J. et al. Neural Circuits for Cognitive Appetite Control in Healthy and Obese Individuals: An fMRI Study. Plos One 10, doi:ARTN e0116640 10.1371/journal.pone.0116640 (2015).

81. Marsh, R. et al. An FMRI study of self-regulatory control and conflict resolution in adolescents with bulimia nervosa. The American journal of psychiatry 168, 1210–1220, doi:10.1176/appi.ajp.2011.11010094 (2011).

82. Marsh, R. et al. Deficient activity in the neural systems that mediate self-regulatory control in bulimia nervosa. Arch Gen Psychiatry 66, 51–63, doi:10.1001/archgenpsychiatry.2008.504 (2009).

83. Balodis, I. M. et al. Divergent neural substrates of inhibitory control in binge eating disorder relative to other manifestations of obesity. Obesity (Silver Spring) 21, 367–377, doi:10.1002/oby.20068 (2013).

84. Eryilmaz, H. et al. Neural determinants of human goal-directed vs. habitual action control and their relation to trait motivation. Scientific Reports 7, doi:ARTN 6002 10.1038/s41598-017-06284-y (2017).

85. Tschernegg, M. et al. Impulsivity relates to striatal gray matter volumes in humans: evidence from a delay discounting paradigm. Frontiers in Human Neuroscience 9, doi:ARTN 384 10.3389/fnhum.2015.00384 (2015).

86. Tang, Y. Y., Posner, M. I., Rothbart, M. K. & Volkow, N. D. Circuitry of self-control and its role in reducing addiction. Trends in Cognitive Sciences 19, 439–444, doi:10.1016/j.tics.2015.06.007 (2015).

87. Lieberz, J. et al. Kinetics of oxytocin effects on amygdala and striatal reactivity vary between women and men. Neuropsychopharmacology, doi:10.1038/s41386-019-0582-6 (2019).

88. Xu, X. et al. Oxytocin biases men but not women to restore social connections with individuals who socially exclude them. Sci Rep 7, 40589, doi:10.1038/srep40589 (2017).

89. Hoge, E. A. et al. Gender moderates the effect of oxytocin on social judgments. Human Psychopharmacology-Clinical and Experimental 29, 299–304, doi:10.1002/hup.2402 (2014).

90. Gao, S. et al. Oxytocin, the peptide that bonds the sexes also divides them. Proc Natl Acad Sci U S A 113, 7650–7654, doi:10.1073/pnas.1602620113 (2016).

91. Feng, C. et al. Oxytocin and vasopressin effects on the neural response to social cooperation are modulated by sex in humans. Brain Imaging Behav 9, 754–764, doi:10.1007/s11682-014-9333-9 (2015).

92. Spengler, F. B. et al. Kinetics and Dose Dependency of Intranasal Oxytocin Effects on Amygdala Reactivity. Biological Psychiatry 82, 885–894, doi:10.1016/j.biopsych.2017.04.015 (2017).

93. Meule, A. et al. Food cue-induced craving in individuals with bulimia nervosa and binge-eating disorder. PLoS One 13, e0204151, doi:10.1371/journal.pone.0204151 (2018).

94. Collins, B. et al. The impact of acute stress on the neural processing of food cues in bulimia nervosa: Replication in two samples. J Abnorm Psychol 126, 540–551, doi:10.1037/abn0000242 (2017).

