## Supplementary material for "Investigating resting brain perfusion abnormalities and target-engagement by intranasal oxytocin in women with bulimia nervosa and binge-eating disorder and healthy controls"

**Methods**

**Additional information regarding participants’ inclusion and exclusion criteria**

Participants were required to be women between 18 and 40 years old, proficient in English, and right handed. Exclusion criteria included pregnancy (tested using a quick urine strip test during the screening session and each experimental session), severe comorbidity (e.g., substance abuse, drug addiction, psychosis, diabetes), history of drug dependence, history of a neurological condition (e.g., epilepsy), a significant visual impairment, which is not corrected by eyewear, currently suffering from a cold or flu, currently smoking > 5 cigarettes per day (past 6 months), consuming > 21 units of alcohol per week, contraindication to MRI scans, and current intake of medication that might potentially interact with oxytocin (e.g., Prostaglandins).

**Clinical characteristics of the participants with BN/BED**

Of the 25 women with BN/BED, 7 women had a comorbid psychiatric disorder. Specifically, 5 women had comorbid depression, 4 women had comorbid generalised anxiety disorder, 4 women had borderline personality disorder, 1 woman had social anxiety, 1 woman had obsessive-compulsive disorder, and 1 woman had an autism spectrum disorder. At the time of the study, 7 women were taking an antidepressant, 1 woman was taking a mood stabiliser, and one woman was taking an antipsychotic drug. Participants with BN/BED reported an average binge eating frequency of 14.14 episodes over the past 28 days (SD = 9.88). The women with BN endorsed an average frequency of self-induced vomiting equal to 10.40 occasions over the past 28 days (SD = 13.61), an average laxative abuse frequency of 5.13 occasions over the past 28 days (SD = 8.35), an average frequency of “hard exercise intended to control weight or shape” equal to 7.31 occasions over the past 28 days (SD = 8.57), and one participant reported using diuretic pills on 4 occasions over the past 28 days.

**Supplementary information on Statistical analysis**

### Fig. S1 – Regions-of-interest: Representation in the radiological convention of the regions-of-interest used in our hypothesis-driven analyses; VTA – Ventral tegmental area; SN – Substantia nigra; Amy – Amygdala; PFC – Medial Prefrontal Cortext; HPT – Hypothalamus; ACG – Anterior Cingulate Gyrus; Acc – Accumbens).

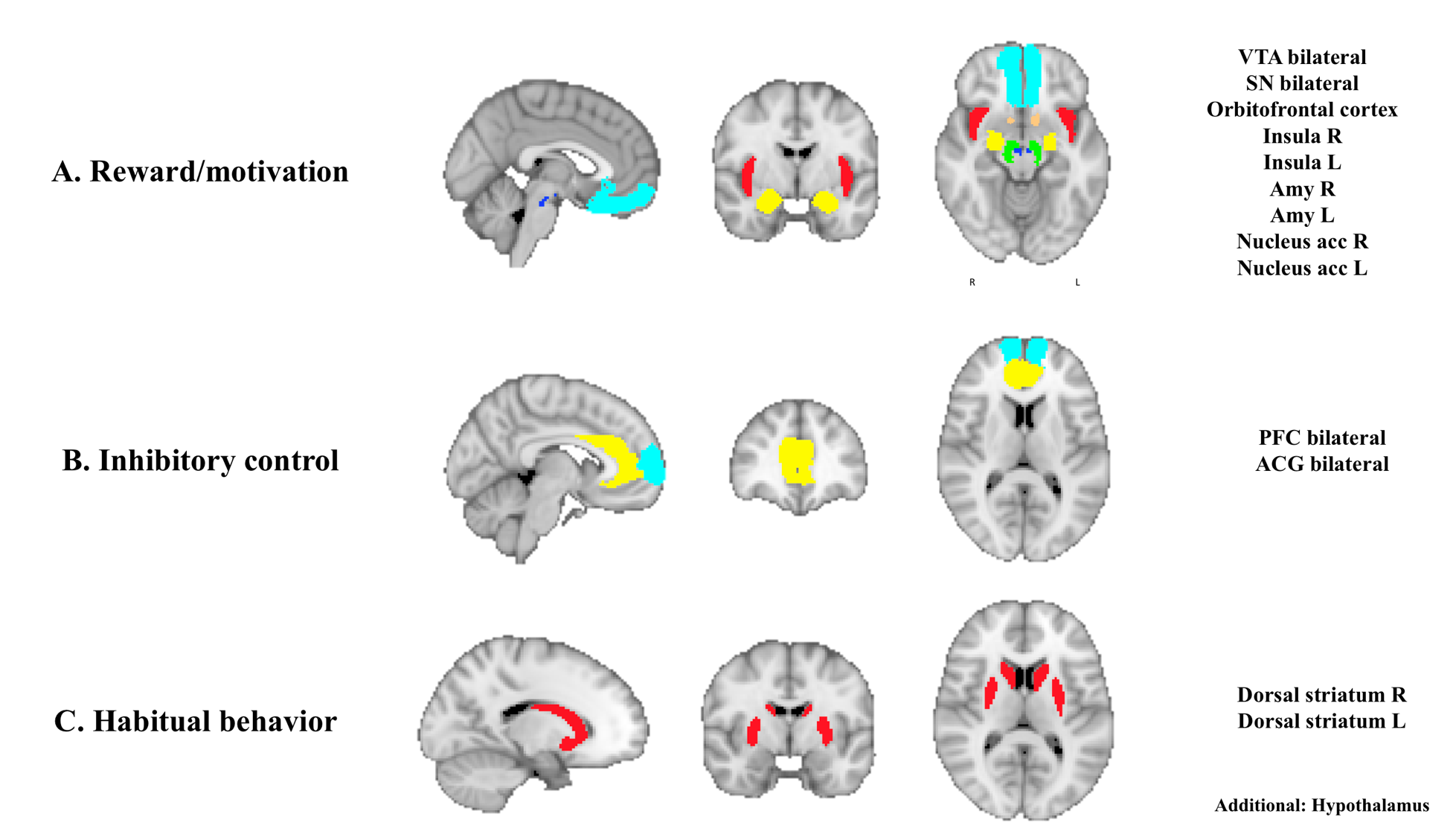

**Whole-brain analyses**

In all of our whole-brain analyses, we took a partitioned errors approach (1) instead of a simple mixed analysis of covariance. This decision was taken to avoid violations of the assumptions of the latest, such as violations of sphericity likely to exist in repeated measures designs (1). For the main effect of diagnosis, we averaged CBF maps across treatment sessions for each subject. We then examined diagnosis effects in independent T-tests across the whole brain. For analyses testing the main effect of treatment, we calculated difference CBF maps resulting from the subtraction between CBF maps acquired in the OT and placebo sessions. We then tested for treatment effects using a one-sample T-test across the whole brain. For the diagnosis x treatment interaction effect, we used independent T-tests to compare subtracted OT-placebo CBF maps between healthy women and BN/BED patients. For all of our whole-brain analyses, we first ran an F-contrast to identify changes regardless of direction of effect. This analysis was then followed by directed T-contrasts to test whether these changes would correspond to increases or decreases in resting rCBF.

**Voxel-based morphometry analysis**

We preprocessed all high-resolution structural images using the Voxel-based Morphometry analysis toolbox CAT12 running on Matlab R2016a (Mathworks Inc., Natick, MA, USA). First, we used the module “Segment Data” of CAT12 to segment each image into grey matter (GM), white matter, and cerebrospinal fluid probability maps. The modulated warped GM images were then normalized to MNI-152 standard space with an isotropic voxel resolution of 2 mm × 2 mm × 2 mm. We then averaged the images of all participants to generate a study population-specific template. Next, using “Display one slice for all images,” we checked the data quality to determine whether reasonable results were obtained by the segmentation and normalization procedures (i.e., if the native volume had artefacts or a wrong orientation). Using a boxplot and correlation matrices, we also checked sample homogeneity to identify outliers by visualizing the correlation between the volumes. The modulated GM map of each individual was smoothed with an 8-mm full width at the half-maximum Gaussian kernel. Finally, using the “Estimate TIV” module, we estimated the total intracranial volume (TIV) for all the subjects.

We then used SPM 12 to examine to compare grey matter volume (GMV) between BN/BED and healthy women (diagnosis effect) in an independent T-test across the whole brain, using TIV as covariate. We first ran an F-contrast to identify changes regardless of direction. This analysis was then followed by directed T-contrasts to test whether these changes would correspond to increases or decreases in GMV. For all of our whole-brain analyses, whole-brain cluster-level inference was applied at α = 0.05 using familywise error (FWE) correction for multiple comparisons and a cluster-forming threshold of *p* = 0.005 (uncorrected).

**Results**

**Fig. S2: Effects of diagnosis, treatment and diagnosis x treatment on global grey matter CBF in BN/BED and healthy women.** Individual values, box plots and violin plots depicting global CBF for each diagnosis/treatment groups; middle horizontal lines represent the median; boxes indicate the 25^th^ and 75^th^ percentiles. We tested the effects of diagnosis, treatment and diagnosis x treatment in mixed analysis of variance. Statistical significance was set to *p* < 0.05; GM – Grey-matter; CBF – Cerebral blood flow.

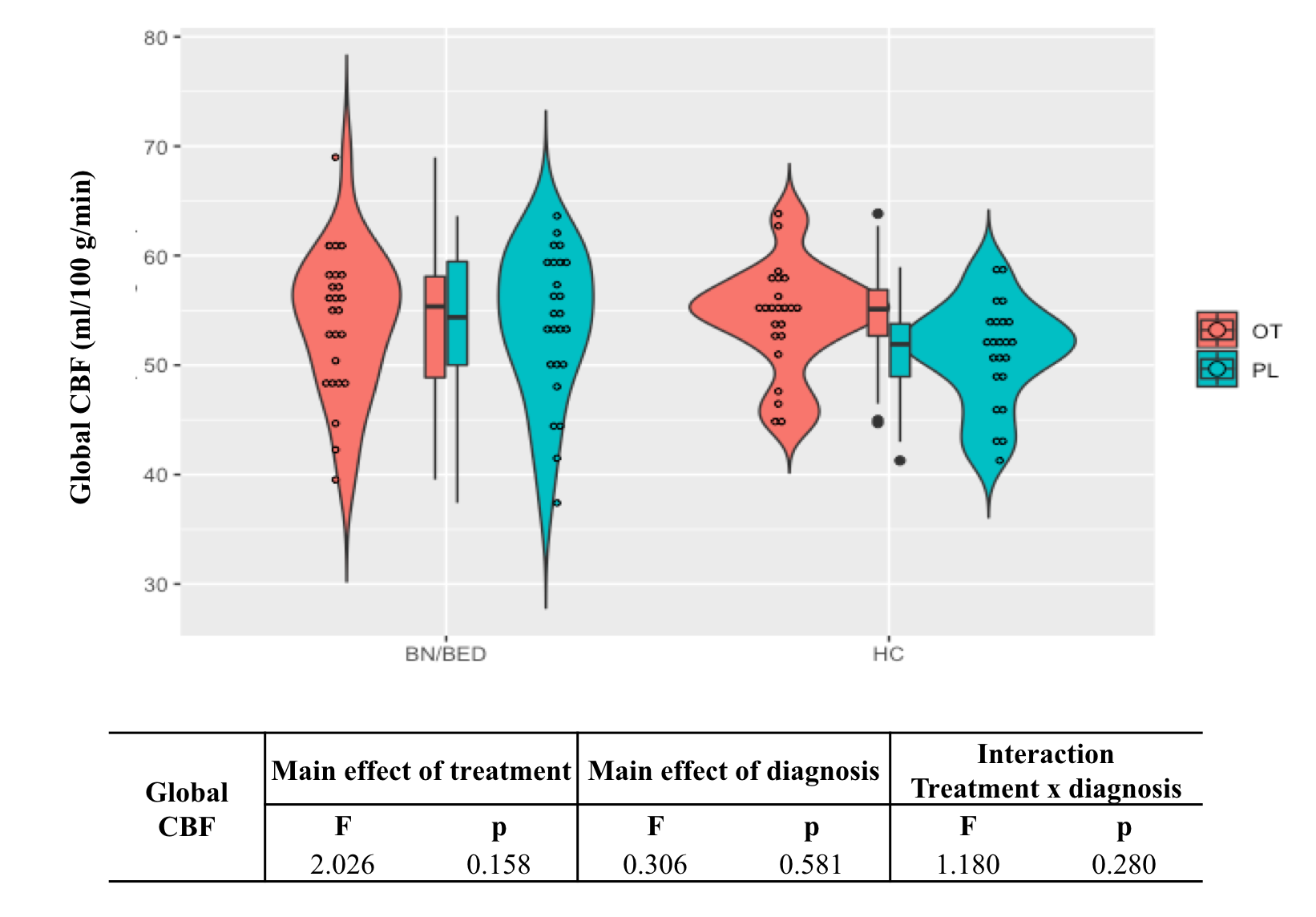

**Table S1 - Effects of diagnosis, treatment and diagnosis x treatment on resting regional cerebral blood flow (rCBF) within neural circuits relevant for BN/BED (accounting for global CBF only).** This table shows the results of a hypothesis-driven investigation of the effects of diagnosis, treatment and diagnosis x treatment on rCBF within 14 anatomical regions-of-interest suggested to be involved in BN/BED. The effects of diagnosis, treatment and diagnosis x treatment were tested in a full factorial linear mixed model, controlling for global grey-matter cerebral blood flow. Statistical significance was set to *p* < 0.05, after correction for multiple testing with the Benjamini-Hochberg procedure; VTA – Ventral tegmental area; SN – Substantia nigra; Amy – Amygdala; PFC – Medial Prefrontal Cortext; HPT – Hypothalamus; ACG – Anterior Cingulate Gyrus; Acc – Accumbens.

| **Region-of-interest** | **Main effect of treatment** | | | **Main effect of diagnosis** | | | **Interaction Treatment x diagnosis** | | |
| --- | --- | --- | --- | --- | --- | --- | --- | --- | --- |
|  | **F** | **P (uncorrected)** | **P (adjusted)** | **F** | **P (uncorrected)** | **P (adjusted)** | **F** | **P (uncorrected)** | **P (adjusted)** |
| VTA | 0.674 | 0.414 | 1.449 | 0.502 | 0.481 | 0.842 | 0.007 | 0.933 | 1.005 |
| SN | 1.244 | 0.268 | 1.876 | 0.011 | 0.917 | 0.988 | 0.020 | 0.887 | 1.129 |
| Right Amy | 0.343 | 0.560 | 1.568 | 1.606 | 0.208 | 0.582 | 0.010 | 0.921 | 1.075 |
| Left amy | 0.258 | 0.613 | 1.430 | 0.440 | 0.509 | 0.792 | 0.135 | 0.714 | 1.249 |
| **PFC** | 1.695 | 0.196 | 2.744 | 5.890 | **0.017** | 0.119 | 0.056 | 0.184 | 2.576 |
| **Orbitofrontal** | 0.025 | 0.874 | 1.359 | 6.002 | **0.016** | 0.224 | 0.046 | 0.830 | 1.291 |
| Insula right | 1x10^-6^ | 0.999 | 0.999 | 3.577 | 0.062 | 0.217 | 0.532 | 0.468 | 1.638 |
| Insula left | 2.6x10^-4^ | 0.987 | 1.063 | 0.218 | 0.641 | 0.748 | 0.001 | 0.974 | 0.974 |
| HPT | 1.098 | 0.297 | 1.386 | 0.798 | 0.374 | 0.873 | 0.035 | 0.852 | 1.193 |
| ACG | 0.013 | 0.908 | 1.271 | 3.788 | 0.055 | 0.257 | 0.272 | 0.603 | 1.206 |
| Dorsal striatum right | 0.002 | 0.968 | 1.129 | 0.687 | 0.409 | 0.818 | 0.527 | 0.470 | 1.316 |
| Dorsal striatum left | 0.004 | 0.951 | 1.210 | 0.264 | 0.608 | 0.851 | 0.669 | 0.416 | 1.941 |
| Acc right | 0.115 | 0.736 | 1.288 | 0.003 | 0.956 | 0.956 | 0.474 | 0.493 | 1.150 |
| Acc left | 0.137 | 0.713 | 1.426 | 0.229 | 0.633 | 0.806 | 0.821 | 0.367 | 2.569 |

**Table S2 - Independent T-contrast comparing resting regional cerebral blood flow (rCBF) in women with BN/BED versus healthy women (accounting for global CBF).** We compared CBF maps from BN/BED women versus healthy women using T-contrasts (to capture the direction of potential rCBF changes), controlling for global CBF as a nuisance variable. We conducted cluster-level inference, reporting clusters significant at p<0.05 FWE-corrected (cluster-forming threshold: p<0.005, uncorrected).

| Cluster Description | Hemisphere | K | P_FWE_ | Peak Coordinates | | | Description |
| --- | --- | --- | --- | --- | --- | --- | --- |
|  |  |  |  | **x** | **y** | **z** |  |
| BN/BED > HC | | | | | | | |
| Cluster 1: Inferior, middle, superior temporal gyri | Right | 2551 | <0.001 | 58 | -32 | -18 | Right middle temporal gyrus |
|  |  |  |  | 54 | -10 | -32 | Right inferior temporal gyrus |
|  |  |  |  | 62 | -20 | -22 | Right middle temporal gyrus |
| Cluster 2: Inferior, middle, superior temporal gyri, planum temporale | Left | 1405 | <0.001 | -54 | -46 | 6 | Left middle temporal gyrus |
|  |  |  |  | -58 | -28 | -22 | Left inferior temporal gyrus |
|  |  |  |  | -56 | -38 | 20 | Left planum temporale |
| Cluster 3: Anterior Cingulate, Superior Frontal gyrus, Medial orbitofrontal cortex, Gyrus Rectus | Bilateral | 593 | <0.001 | -6 | 56 | 2 | Left superior frontal gyrus |
|  |  |  |  | -4 | 56 | 14 | Left superior frontal gyrus |
|  |  |  |  | -4 | 60 | -14 | Left gyrus rectus |

**Table S3 - Independent T-contrast comparing resting regional cerebral blood flow (rCBF) in women with BN/BED versus healthy women (accounting for global CBF and BMI).** We compared CBF maps from BN/BED women versus healthy women using T-contrasts (to capture the direction of potential rCBF changes), controlling for global CBF and BMI as nuisance variables. We conducted cluster-level inference, reporting clusters significant at p<0.05 FWE-corrected (cluster-forming threshold: p<0.005, uncorrected).

| Cluster Description | Hemisphere | K | P_FWE_ | Peak Coordinates | | | Description |
| --- | --- | --- | --- | --- | --- | --- | --- |
|  |  |  |  | **x** | **y** | **z** |  |
| BN/BED > HC | | | | | | | |
| Cluster 1: Inferior, middle, superior temporal gyri, posterior insula | Right | 3626 | <0.001 | 54 | -10 | -32 | Right inferior temporal gyrus |
|  |  |  |  | 62 | -20 | -22 | Right middle temporal gyrus |
|  |  |  |  | 60 | -32 | -18 | Right middle temporal gyrus |
| Cluster 2: Inferior, middle, superior temporal gyri, planum temporale | Left | 1640 | <0.001 | -54 | -46 | 6 | Left middle temporal gyrus |
|  |  |  |  | -58 | -52 | -6 | Left middle temporal gyrus |
|  |  |  |  | -56 | -40 | 18 | Left planum temporale |
| Cluster 3: Anterior Cingulate, Superior Frontal gyrus, Medial orbitofrontal cortex, Gyrus Rectus | Bilateral | 2070 | <0.001 | -4 | 58 | 2 | Left superior frontal gyrus |
|  |  |  |  | -4 | 56 | 14 | Left superior frontal gyrus |
|  |  |  |  | 20 | 38 | 34 | Right superior frontal gyrus |

**Table S4 - Increases in resting regional cerebral blood flow (rCBF) in the brain of BN/BED patients correlate positively with eating disorder symptom severity (adjusting for global CBF, BMI and depression, anxiety and stress scores).** This table shows the result of partial Pearson correlations between mean rCBF in each of the four anatomical regions-of-interest where we found significant differences between the BN/BED and healthy groups and eating disorder symptom severity as measured by the global EDEQ scores, for controls and patients separately; Partial Pearson correlations were calculated with bootstrapping (1000 samples), adjusting for global CBF, BMI and the first principal component summarizing stress, anxiety and depression scores. Statistical significance was set to *p* < 0.05 and is highlighted with the symbol *. Due to some missing data, we present in the row N (valid cases) the amount of available data that was used for the calculation of each correlation. In the last column, we present the result of the statistical comparison of the correlations between the two groups, as assessed by Fisher r-to-z transformation. PFC – Medial Prefrontal cortex; OFC – Orbitofrontal cortex; ACG – Anterior Cingulate Gyrus.

|  | | **Global EDEQ** | | |
| --- | --- | --- | --- | --- |
|  | | **BN/BED** | **HC** | **Group comparison** |
| **PFC** | **r (Pearson)** | 0.293 | -0.181 | Z = 1.411 |
|  | **p** | 0.051 | 0.446 | p = 0.079 |
| **OFC** | **r (Pearson)** | 0.348* | 0.230 | Z = 0.375 |
|  | **p** | 0.019 | 0.329 | p = 0.354 |
| **ACG** | **r (Pearson)** | 0.371* | 0.164 | Z = 0.653 |
|  | **p** | 0.012 | 0.489 | p = 0.257 |
| **Right Insula** | **r (Pearson)** | 0.462* | 0.297 | Z = 0.564 |
|  | **p** | 0.001 | 0.203 | p = 0.287 |
| **N (valid cases)** | | **21** | **19** |  |

**Fig. S3 – Increases in resting regional cerebral blood flow (rCBF) in the brain of BN/BED patients (accounting for global CBF only).** This figure shows the results of a directed T-contrast where we tested for increases (BN/BED > Controls) or decreases (Controls > BN/BED) in rCBF in patients compared to controls, accounting for global grey-matter cerebral blood only, at the whole-brain level. Whole-brain cluster-level inference was applied at α = 0.05 using familywise error (FWE) correction for multiple comparisons and a cluster-forming threshold of *p* = 0.005 (uncorrected). Images are shown as T-statistic in radiological convention. We did not find any significant cluster depicting decreases in rCBF in BN/BED patients (compared to healthy controls).

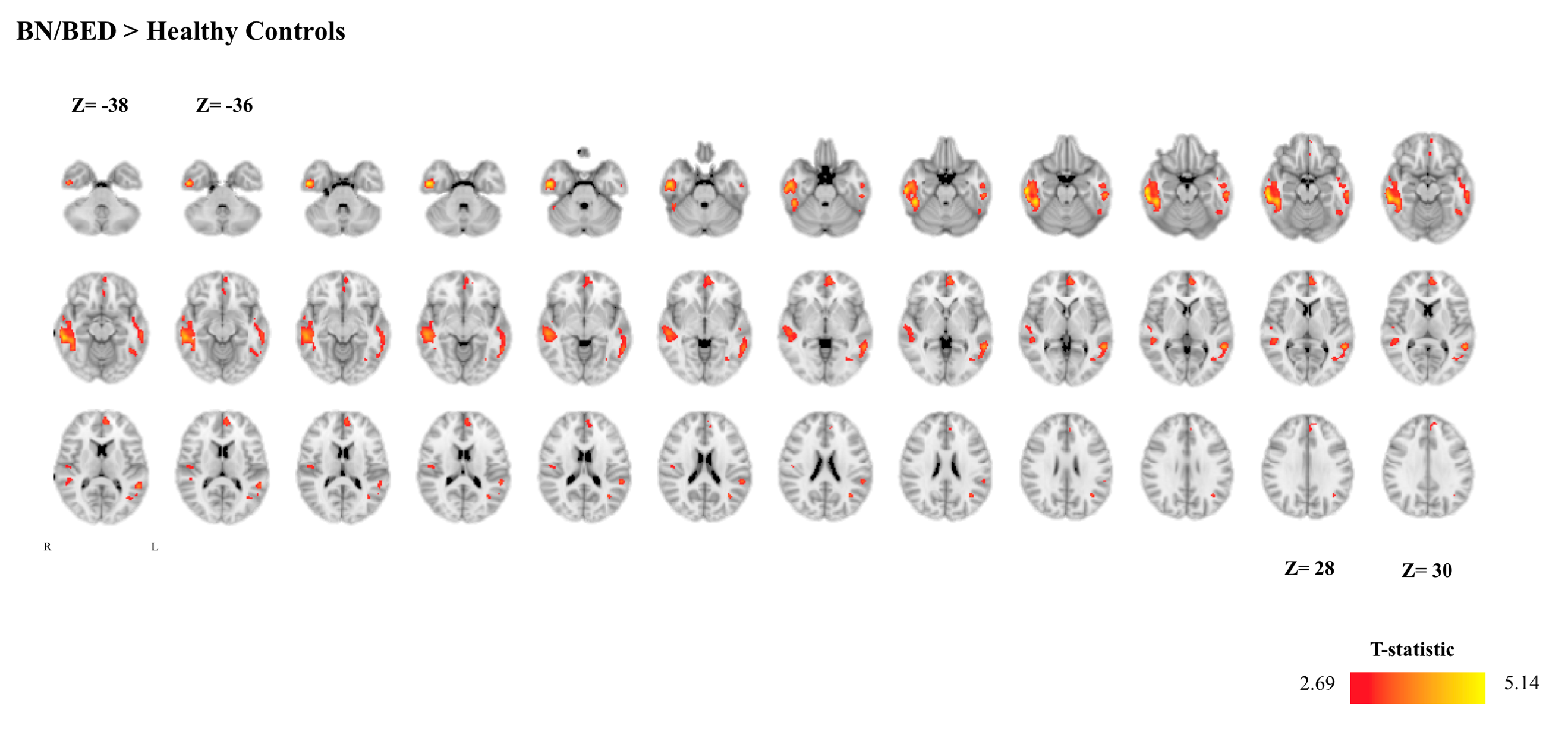

**Table S5 - Effects of diagnosis and treatment on resting regional cerebral blood flow (rCBF) within neural circuits relevant for BN/BED (accounting for hormonal contraception use).** This table shows the results of a hypothesis-driven investigation of the effects of diagnosis, treatment and diagnosis x treatment on rCBF within 14 anatomical regions-of-interest suggested to be involved in BN/BED. We tested these effects in a full factorial linear mixed model, controlling for global grey-matter cerebral blood flow and hormonal contraception use. Statistical significance was set to *p* < 0.05, after correction for multiple testing with the Benjamini-Hochberg procedure. VTA – Ventral tegmental area; SN – Substantia nigra; Amy – Amygdala; PFC – Medial Prefrontal Cortext; HPT – Hypothalamus; ACG – Anterior Cingulate Gyrus; Acc – Accumbens.

| **Region-of-interest** | **Main effect of treatment** | | | **Main effect of diagnosis** | | | **Interaction Treatment x diagnosis** | | |
| --- | --- | --- | --- | --- | --- | --- | --- | --- | --- |
|  | **F** | **P (uncorrected)** | **P (adjusted)** | **F** | **P (uncorrected)** | **P (adjusted)** | **F** | **P (uncorrected)** | **P (adjusted)** |
| VTA | 0.764 | 0.385 | 1.348 | 0.536 | 0.466 | 0.816 | 0.119 | 0.731 | 1.023 |
| SN | 1.158 | 0.285 | 1.995 | 0.015 | 0.901 | 0.901 | 0.137 | 0.712 | 1.246 |
| Right Amy | 0.342 | 0.560 | 1.568 | 1.704 | 0.196 | 0.549 | 0.346 | 0.558 | 2.604 |
| Left amy | 0.175 | 0.676 | 1.183 | 0.364 | 0.548 | 0.852 | 0.137 | 0.712 | 1.108 |
| **PFC** | 1.940 | 0.168 | 2.352 | 5.184 | **0.025** | 0.175 | 0.023 | 0.879 | 0.947 |
| **Orbitofrontal** | 0.177 | 0.675 | 1.350 | 6.083 | **0.016** | 0.224 | 0.082 | 0.776 | 0.988 |
| Insula right | 0.017 | 0.898 | 1.143 | 3.148 | 0.080 | 0.373 | 0.227 | 0.635 | 1.778 |
| Insula left | 0.010 | 0.992 | 0.992 | 0.182 | 0.671 | 0.939 | 0.011 | 0.916 | 0.916 |
| HPT | 0.970 | 0.328 | 1.531 | 1.354 | 0.248 | 0.579 | 0.039 | 0.844 | 0.985 |
| ACG | 0.095 | 0.759 | 1.063 | 1.889 | 0.173 | 0.606 | 0.145 | 0.704 | 1.408 |
| Dorsal striatum right | 0.010 | 0.921 | 1.075 | 0.911 | 0.343 | 0.686 | 0.223 | 0.638 | 1.489 |
| Dorsal striatum left | 0.001 | 0.973 | 1.048 | 0.134 | 0.716 | 0.911 | 1.187 | 0.279 | 3.906 |
| Acc right | 0.186 | 0.667 | 1.556 | 0.029 | 0.865 | 0.931 | 0.249 | 0.619 | 2.167 |
| Acc left | 0.168 | 0.683 | 1.062 | 0.117 | 0.733 | 0.855 | 1.042 | 0.310 | 2.170 |

**Fig. S4 – Increases in resting regional cerebral blood flow (rCBF) in the brain of BN/BED patients (accounting for hormonal contraception use).** This figure shows the results of a directed T-contrast where we tested for increases (BN/BED > Controls) or decreases (Controls > BN/BED) in rCBF in patients compared to controls at the whole-brain level, accounting for global grey-matter cerebral blood flow and hormonal contraception use. Whole-brain cluster-level inference was applied at α = 0.05 using familywise error (FWE) correction for multiple comparisons and a cluster-forming threshold of *p* = 0.005 (uncorrected). Images are shown as T-statistic in radiological convention. We did not find any significant cluster depicting decreases in rCBF in BN/BED patients (compared to healthy women).

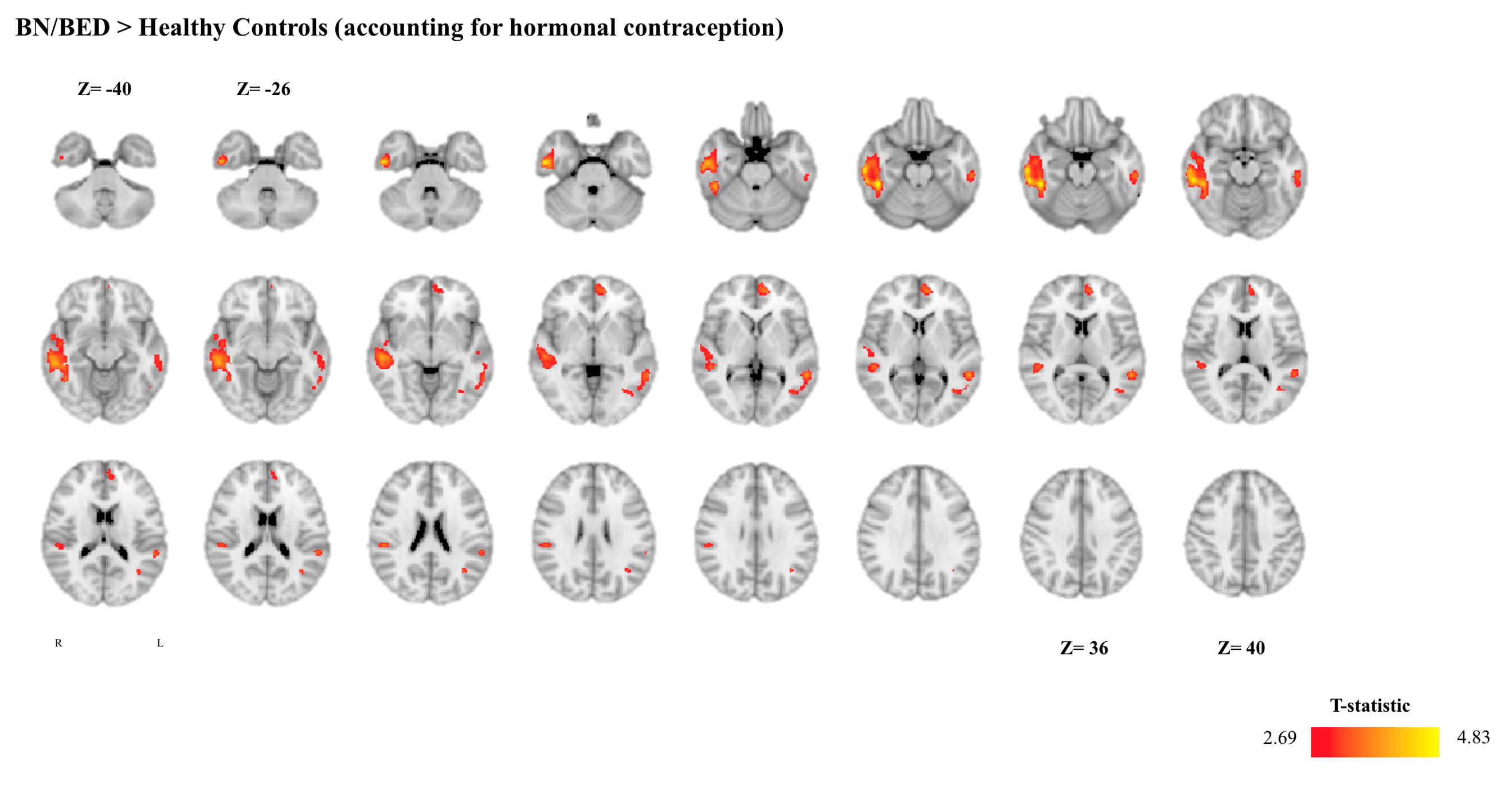

**Fig. S5 – Increases in resting regional cerebral blood flow (rCBF) in the brain of BN/BED patients (excluding BED).** This figure shows the results of a directed T-contrast where we tested for increases (BN/BED > Controls) or decreases (Controls > BN/BED) in rCBF in BN patients (excluding the 5 BED patients) compared to controls at the whole-brain level, accounting for global grey-matter cerebral blood flow. Whole-brain cluster-level inference was applied at α = 0.05 using familywise error (FWE) correction for multiple comparisons and a cluster-forming threshold of *p* = 0.005 (uncorrected). Images are shown as T-statistic in radiological convention. We did not find any significant cluster depicting decreases in rCBF in BN/BED patients (compared to healthy women).

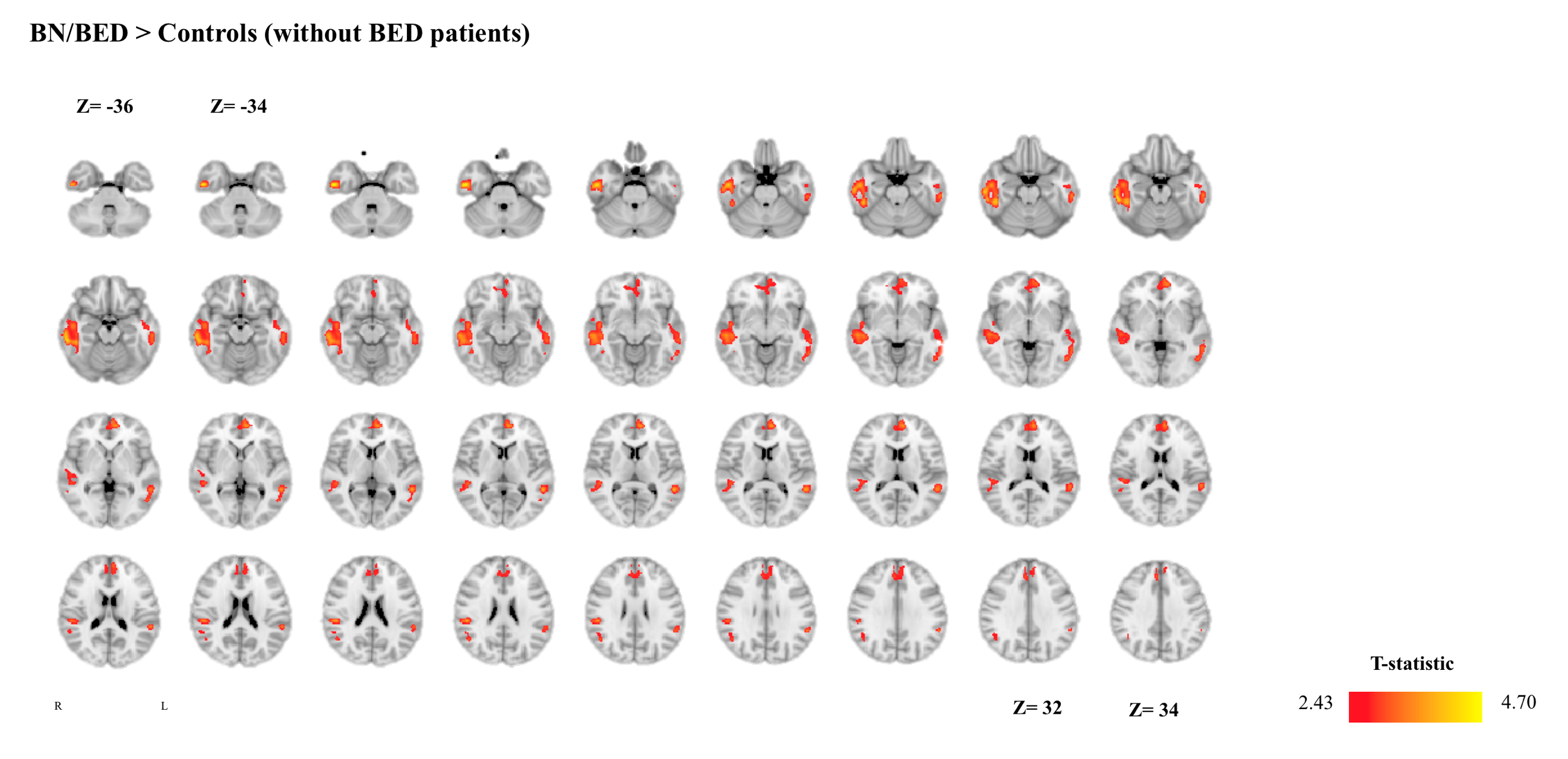

**Fig. S6 – Increases in resting regional cerebral blood flow (rCBF) in the brain of BN/BED patients (accounting for diagnostic category).** This figure shows the results of a directed T-contrast where we tested for increases (BN/BED > Controls) or decreases (Controls > BN/BED) in rCBF in patients compared to controls at the whole-brain level, accounting for global grey-matter cerebral blood flow and for one categorical variable representing diagnostic category: healthy control, BN or BED. Whole-brain cluster-level inference was applied at α = 0.05 using familywise error (FWE) correction for multiple comparisons and a cluster-forming threshold of *p* = 0.005 (uncorrected). Images are shown as T-statistic in radiological convention. We did not find any significant cluster depicting decreases in rCBF in BN/BED patients (compared to healthy women).

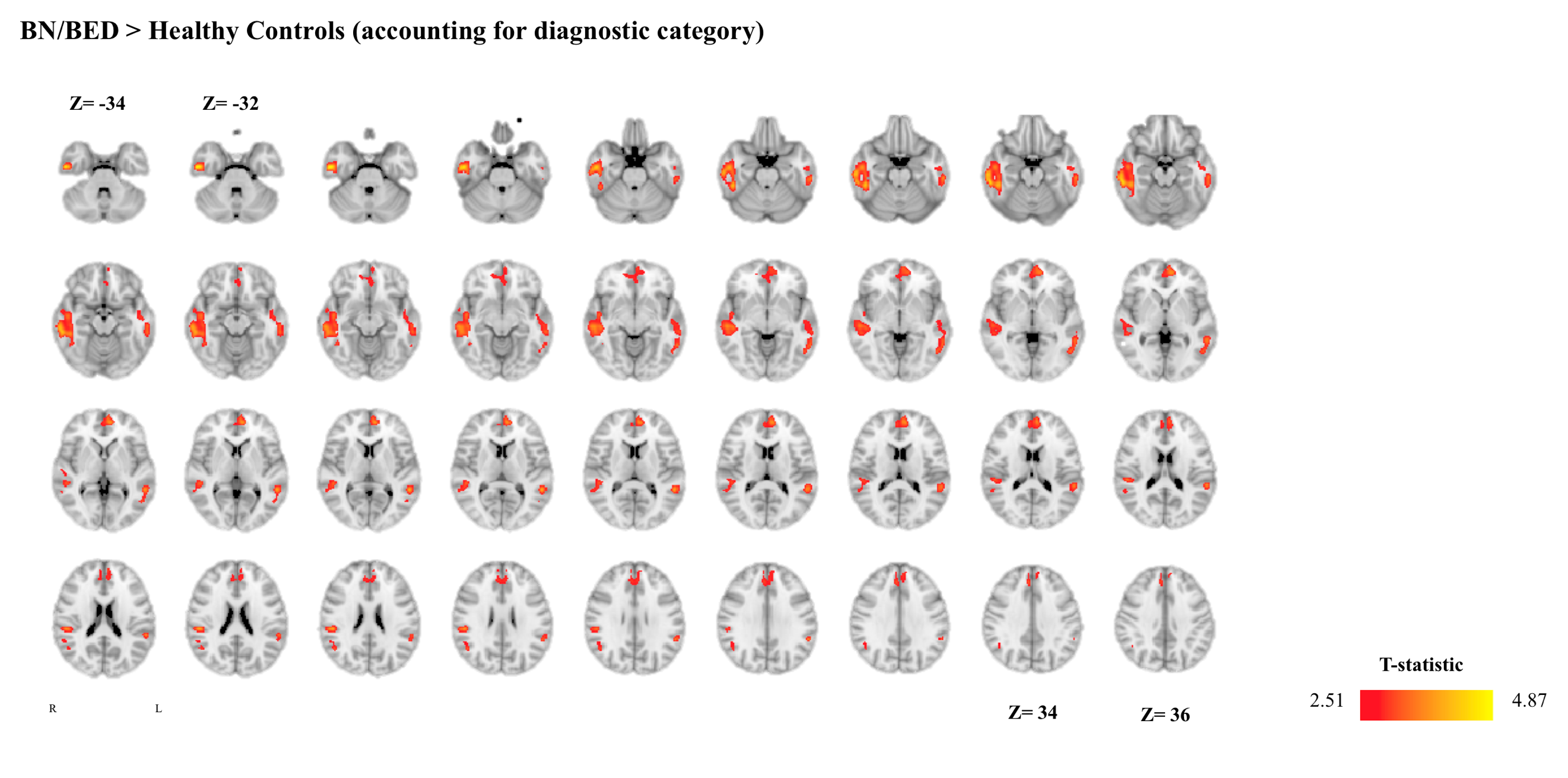

**Fig. S7 – Increases in resting regional cerebral blood flow (rCBF) in the brain of BN/BED patients (accounting for comorbidities/current treatment).** This figure shows the results of a directed T-contrast analysis at the whole-brain level where we tested for increases (BN/BED > Controls) or decreases (Controls > BN/BED) in rCBF in patients compared to controls, accounting for global grey-matter cerebral blood flow and for one categorical binary variable representing the current existence of any comorbidities or pharmacological treatment. Whole-brain cluster-level inference was applied at α = 0.05 using familywise error (FWE) correction for multiple comparisons and a cluster-forming threshold of *p* = 0.005 (uncorrected). Images are shown as T-statistic in radiological convention. We did not find any significant cluster depicting decreases in rCBF in BN/BED patients (compared to healthy women).

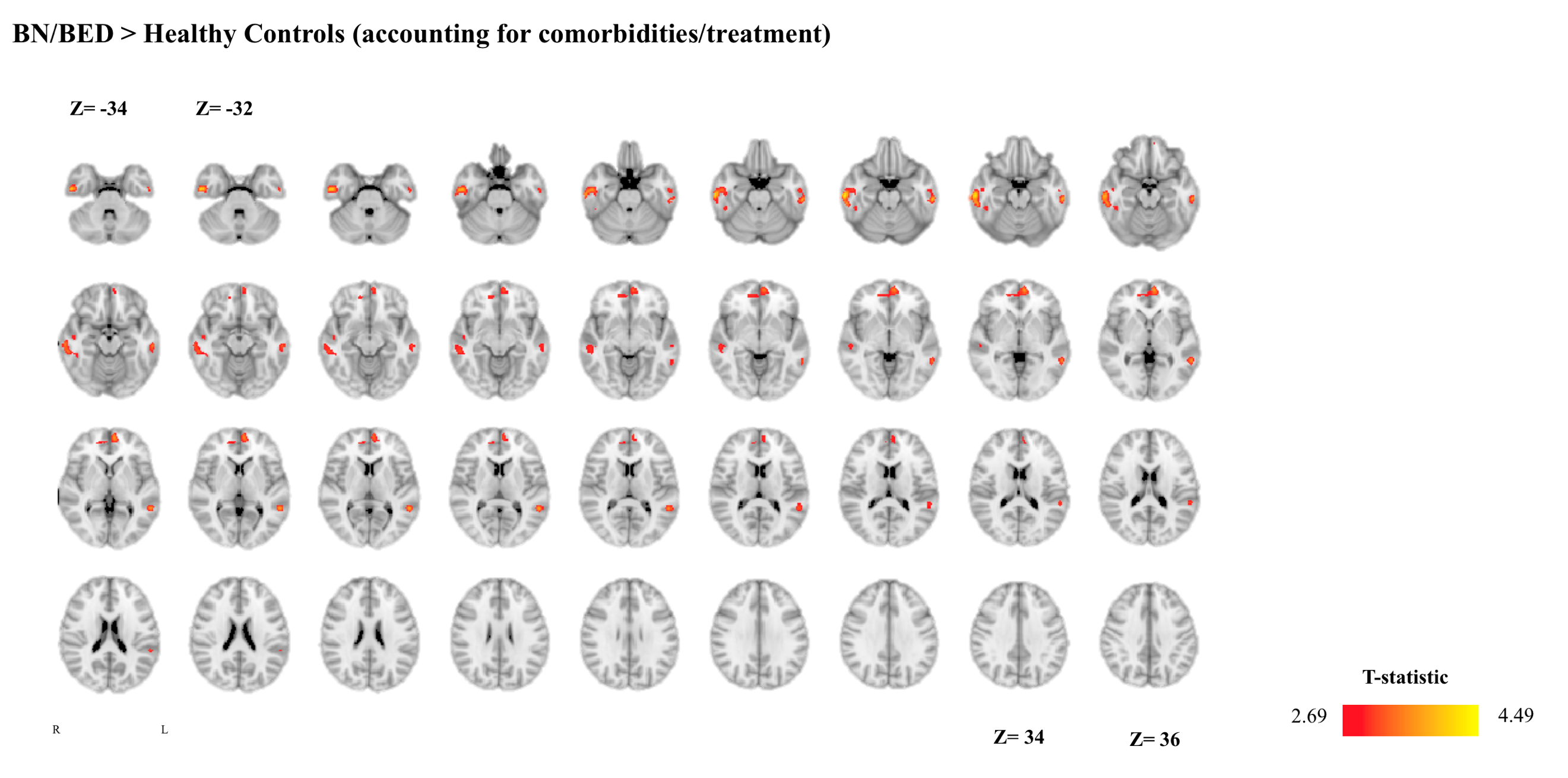

**Fig. S8 – Increases in resting regional cerebral blood flow (rCBF) in the brain of BN/BED patients (accounting for grey-matter volume).** This figure shows the results of a directed T-contrast where we tested for increases (BN/BED > Controls) or decreases (Controls > BN/BED) in rCBF in patients compared to controls at the whole-brain level, accounting for global grey-matter cerebral blood flow and grey-matter volume (GMV) within the clusters where we found patients to have higher rCBF than controls. Whole-brain cluster-level inference was applied at α = 0.05 using familywise error (FWE) correction for multiple comparisons and a cluster-forming threshold of *p* = 0.005 (uncorrected). Images are shown as T-statistic in radiological convention. We did not find any significant cluster depicting decreases in rCBF in BN/BED patients (compared to healthy women).

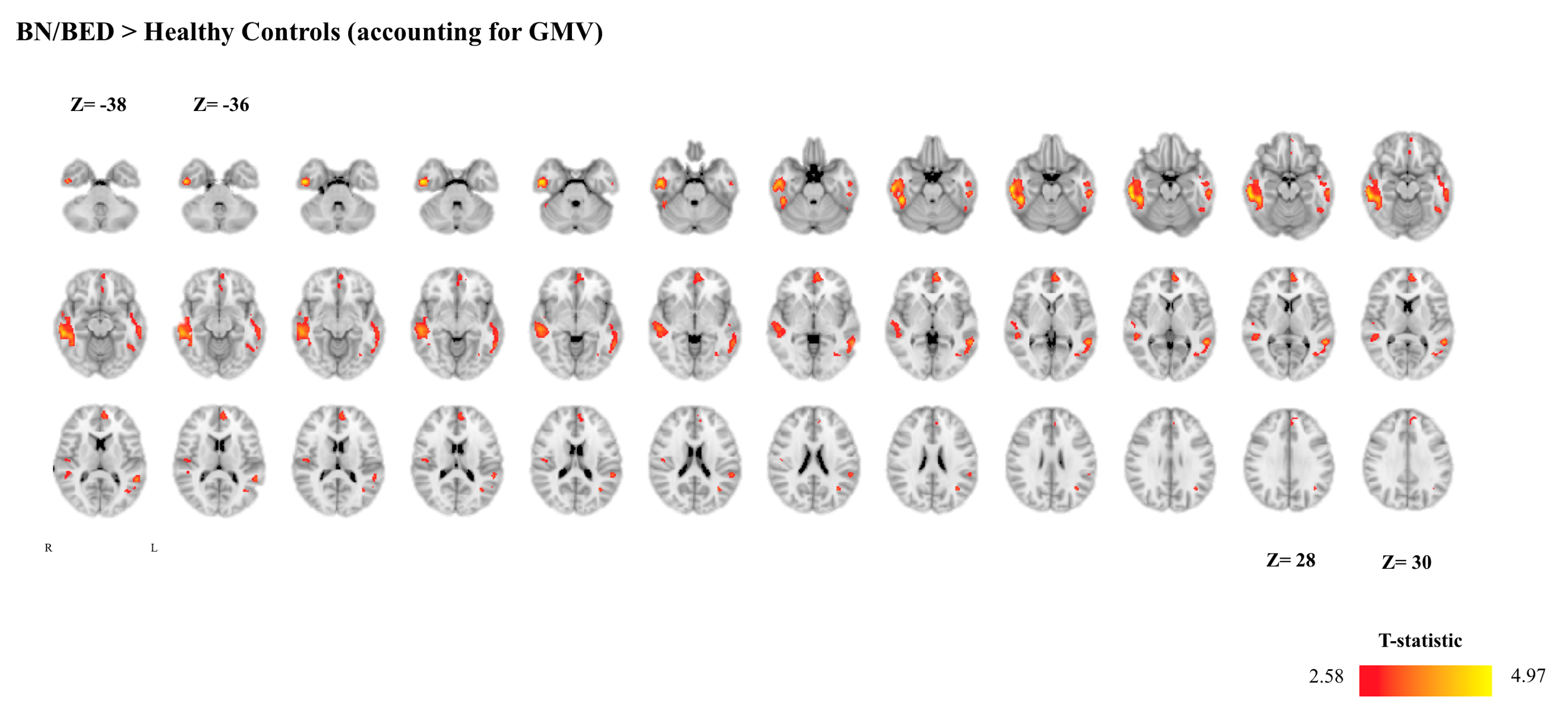

**Fig. S9 – Co-representation of the changes in resting regional cerebral blood flow (rCBF) and grey matter volume (GMV) identified in BN/BED compared to healthy women.** In this figure, we show the spatial distribution of the clusters where we found significant differences between BN/BED and healthy women in rCBF and GMV. Images are shown as T-statistic in radiological convention. In yellow, we present the clusters where we found BN/BED patients to present higher rCBF than healthy controls. In blue, we present the clusters where we found BN/BED patients to present lower GMV than healthy women. Whole-brain cluster-level inference was applied at α = 0.05 using familywise error (FWE) correction for multiple comparisons and a cluster-forming threshold of *p* = 0.005 (uncorrected). Images are shown as T-statistic in radiological convention. The circle highlights an area where BN/BED patients presented both a decrease in GMV and an increase in rCBF in the right temporal lobe. We did not find any significant cluster where BN/BED patients presented higher GMV than healthy controls.

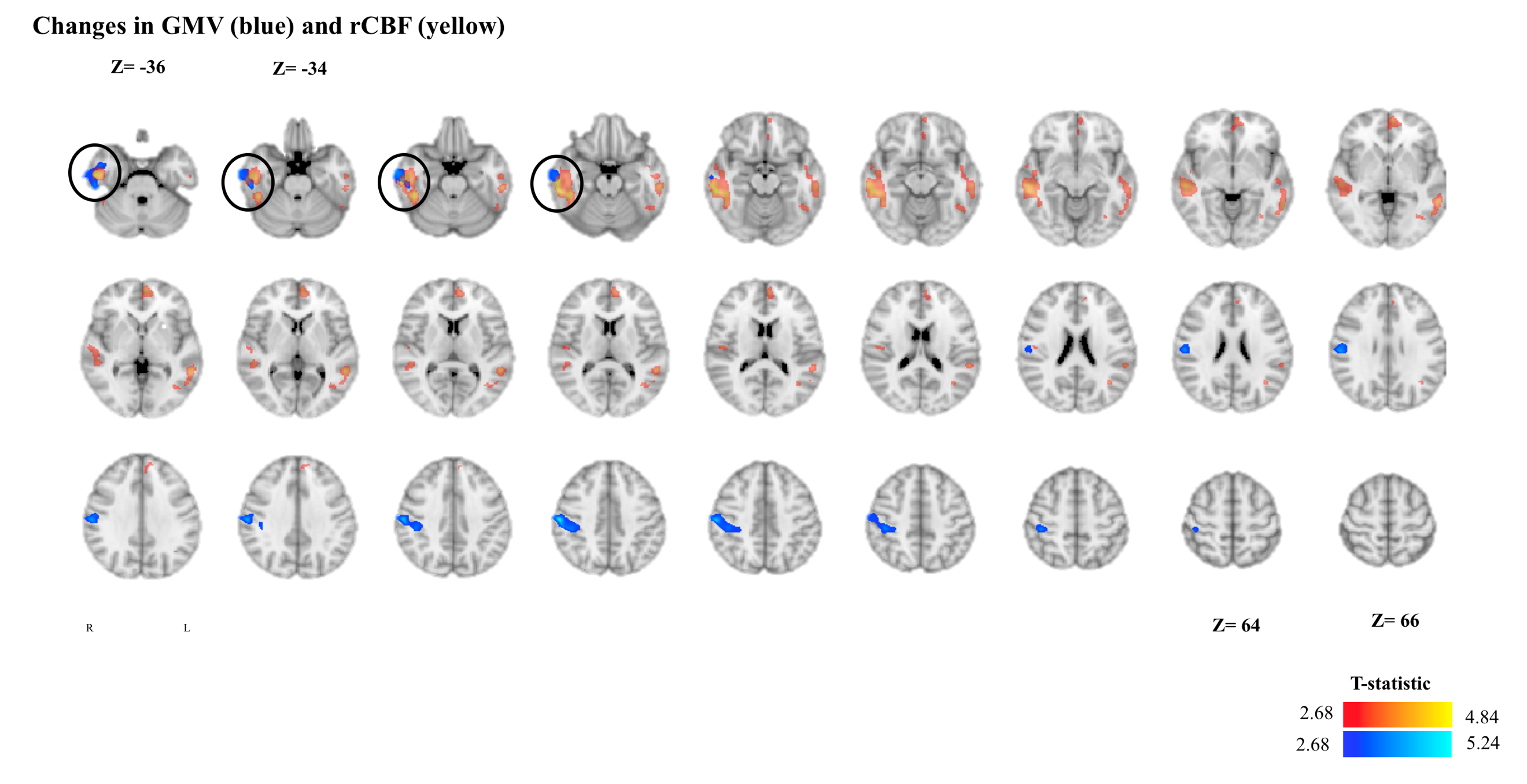

**Fig. S10 – Grey matter volume (GMV) correlates negatively with resting regional cerebral blood flow (rCBF) in the right temporal lobe of BN/BED patients.** In this figure, we show the results of a Pearson correlation analysis (with bootstrapping 1000 samples) examining the association between estimates of GMV and rCBF in the right temporal lobe of BN/BED patients. We found a significant negative correlation between GMV and rCBF in this area for BN/BED patients, but not controls. The scatter plot depicts the association between these two variables for BN/BED (blue) and healthy (yellow) women separately.

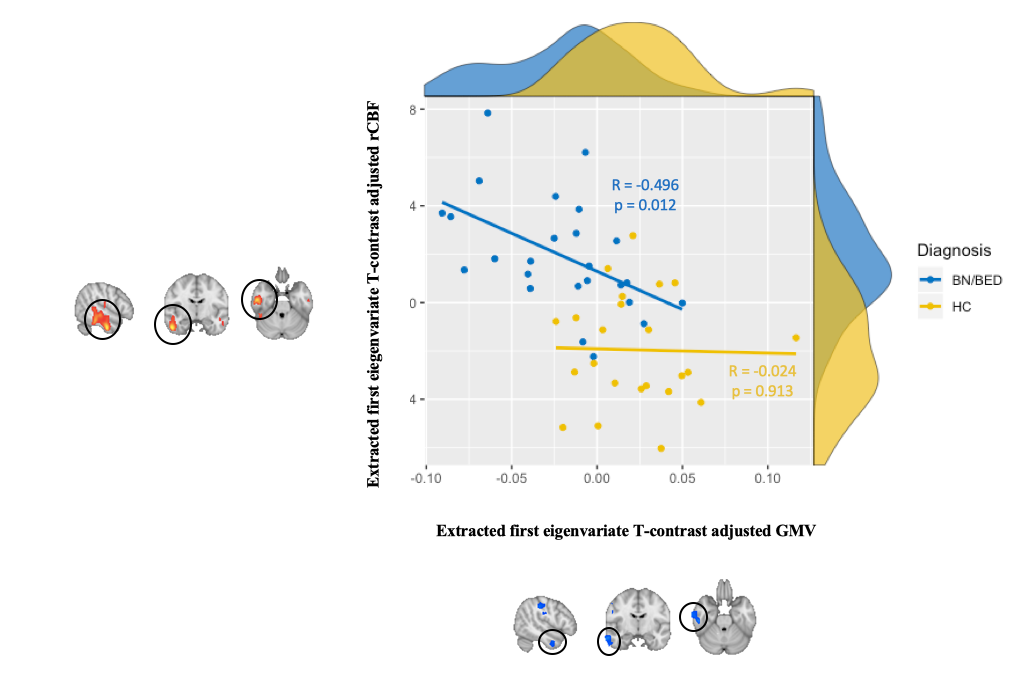

**Supplementary References**

1.	McFarquhar M, McKie S, Emsley R, Suckling J, Elliott R, Williams S. Multivariate and repeated measures (MRM): A new toolbox for dependent and multimodal group-level neuroimaging data (vol 132, pg 373, 2016). Neuroimage. 2016;137:213-.
